# Optimal investments in private land conservation depend more on landholder preferences than climate change

**DOI:** 10.1101/2023.11.26.568746

**Authors:** Brooke A. Williams, Carla L. Archibald, James Brazill-Boast, Michael J. Drielsma, Rajesh Thapa, Jamie Love, Frankie H. T. Cho, Daniel Lunney, James A. Fitzsimons, Sayed Iftekhar, Jaramar Villarreal-Rosas, Sarah Bekessy, Scott Benitez Hetherington, Clive A. McAlpine, Linda Beaumont, Jillian Thonell, Jonathan R. Rhodes

**Affiliations:** School of the Environment, The University of Queensland, Brisbane, QLD, 4072, Australia; Centre for Biodiversity and Conservation Science, The University of Queensland, Brisbane, QLD, 4072, Australia; School of Life and Environmental Sciences, Deakin University, 221 Burwood Highway, Burwood, VIC, 3125, Australia; Department of Planning and Environment, Locked Bag 5022, Parramatta, NSW, 2124, Australia; School of Environmental and Rural Science, The University of New England, Armidale, NSW, 2351, Australia; Land, Environment, Economics and Policy Institute, Department of Economics, University of Exeter, United Kingdom, EX44PU; Faculty of Science, School of Life and Environmental Sciences, University of Sydney, Sydney, NSW 2006, Australia; Australian Museum, 1 William Street, Sydney, NSW 2010, Australia; The Nature Conservancy, Suite 2-01, 60 Leicester Street, Carlton VIC 3053, Australia; School of Law, University of Tasmania, Private Bag 89, Hobart TAS 7001, Australia; Department of Accounting, Finance and Economics, Griffith University, Brisbane, QLD, Australia; Australian Rivers Institute, Griffith University, 170 Kessels Road, Nathan, QLD, 4111, Australia; School of Global, Urban, and Social Studies, RMIT University, Melbourne, VIC, 3000, Australia; Tweed Shire Council, 10-14 Tumbulgum Road, Murwillumbah, NSW, 2484, Australia; School of Natural Sciences, Macquarie University, Sydney, NSW, 2109, Australia

**Keywords:** Conservation tender, landholder behaviour, willingness to accept, koala, Australia, conservation covenant, private protected areas, systematic conservation planning, conservation easement, willingness to participate

## Abstract

Effective private land conservation strategies that consider both landholder preferences and future climatic conditions are critical for preserving biodiversity and ecosystem services. Yet, the interaction and relative importance of these factors for conservation planning performance is unknown. Here, we assess the importance of considering landholder preferences and climate change for prioritising locations for conservation tenders to recruit landholders for conservation covenants. To achieve this we develop a planning framework that accounts for the tender process to optimise investment across regions and apply it to koala-focused tenders in New South Wales (NSW), Australia, exploring four planning approaches that consider or are ignorant to landholder preferences and the tender process and/or climate change. We find that optimal investments depend more on landholder preferences than climate change, and when landholder preferences are ignored, there is little benefit in accounting for climate change. Our analysis reveals new insights into this important interaction.

## Introduction

Nations have expanded their protected area networks (Maxwell et al. 2020), and while protected areas are important for biodiversity conservation, it is increasingly recognised that threats must also be managed beyond protected area boundaries (Kearney et al. 2022). Conservation on private land has therefore become important for biodiversity conservation with researchers working to incorporate landholder values, interests, and priorities into the design of private land conservation programs and planning frameworks to guide their implementation (Cortés Capano et al. 2019). However, incorporating these factors is complicated by the impact of climate change, which is predicted to become a more prominent cause of species declines over the coming century (IPCC 2023). Effective private land conservation strategies that account for the interaction between future climatic conditions and landholder preferences are therefore critical to national and regional plans for preserving biodiversity and ecosystem services. Yet, while incorporating climate change into conservation planning has been a recent area of active research (Jones et al. 2016), the relative importance of climate change and landholder preferences for conservation planning performance is unknown.

A wide range of private land conservation instruments are used to achieve biodiversity conservation with different levels of legal protection and timeframes, yet in many countries conservation covenants (or conservation easements) are widely used to protect biodiversity on private land (Kamal et al. 2015). These are attractive because conservation covenants are legally binding agreements, registered on the property title, between an authorised organisation such as the government and a landholder and provide relatively strong protection (Fitzsimons & Carr 2014). In Australia, for example, conservation covenants are considered to be the highest level of protection on private land as they have strong legal protections compared to other agreement types and are usually in perpetuity (Fitzsimons & Carr 2014).

Successfully establishing conservation covenants relies on landholders’ willingness to participate in covenanting programs, and these are driven by complex socio-economic factors (Kamal et al. 2015). Many factors such as duration of land ownership, knowledge of conservation programs, economics, and how well the program aligns with landowner values can influence program participation (Selinske et al. 2022). Importantly, these factors are highly heterogeneous across any given landscape, as well as being dynamic through time (Nielsen et al. 2017) and it is increasingly recognised that understanding a landholder’s willingness and capacity to participate is crucial for the effective design of private land conservation (Gooden & ’t Sas-Rolfes 2020). In addition, many ecological values on private land targeted by conservation programs are threatened by climate change (Gooden & ’t Sas-Rolfes 2020). Climate change complicates private land conservation efforts by causing species distributions to shift, increasing extinction risks, and leading to large-scale range reductions (Adams-Hosking et al. 2011, 2012; Graham et al. 2019). Therefore, two key considerations when deciding where to invest resources for conservation covenants are spatial variation in landholder preferences and opportunities for uptake in conjunction with the impact of climate change.

Conservation tenders (also known as reverse auctions or procurement auctions) are a common way to recruit landholders for conservation covenants by providing a mechanism for landholders to submit bids on the financial payment they would require to have a covenant on their property (Whitten et al. 2017). Tenders are often used over fixed-rate financial payment schemes as it theoretically improves cost effectiveness because landholders are able to best estimate costs in relation to their own enterprises, auction prices are more likely to reflect the true cost of the resources being used to produce the environmental outcome, and the scope for rent seeking is reduced (Rolfe et al. 2022). Yet, deciding where to invest in the delivery of conservation tenders is a significant challenge as decision makers must consider ecological, climate and socio-economic factors (Fitzsimons & Cooke 2021). Ignoring these factors will likely result in suboptimal decisions, but how suboptimal is unknown.

Here, we assess the implications of ignoring landholder preferences and future impacts of climate change for prioritising locations to deliver tenders for recruiting landholders to a conservation covenant program. To achieve this, we develop a novel spatial planning framework that accounts for the entire conservation tender process to optimise investment across different geographic regions, while considering landholder preferences and future climate conditions. We use New South Wales (NSW), Australia as a case study and identify priority locations for private land conservation through tender investment which focus on conserving habitat for the koala (*Phascolarctos cinereus*), an endangered species in the state that is a priority for conservation (Blanch et al. 2022; Department of the Environment 2022). Private land conservation is particularly important for koala populations as 77% of their range is located on private lands (Kearney et al. 2022). By comparing conservation outcomes when landholder preferences and climate change are considered or are not considered we explicitly quantify the implications of these two factors. In doing so, we provide important insight into their interaction while providing an evidence base to inform planning for conservation tender investment in the region.

## Methods

### Study region

Our study region encompasses the North Coast, Northern Tablelands, Northwest Slopes, Central Coast, the Central and Southern Tablelands, and the South Coast regions of NSW (Figure 1). These regions cover most of the koala’s range in NSW and are priority areas for koala conservation in the state. To reflect the unit of management at which agencies deliver private land conservation and tenders (in NSW this is The Biodiversity Conservation Trust (BCT), a statutory not-for-profit body (Government of New South Wales 2016; Biodiversity Conservation Trust 2018)) we selected Local Government Areas (LGAs) as the spatial planning unit (n = 102), which contain properties considered suitable for private land conservation (n = 199,416; see SM1.1).

**Figure 1.**
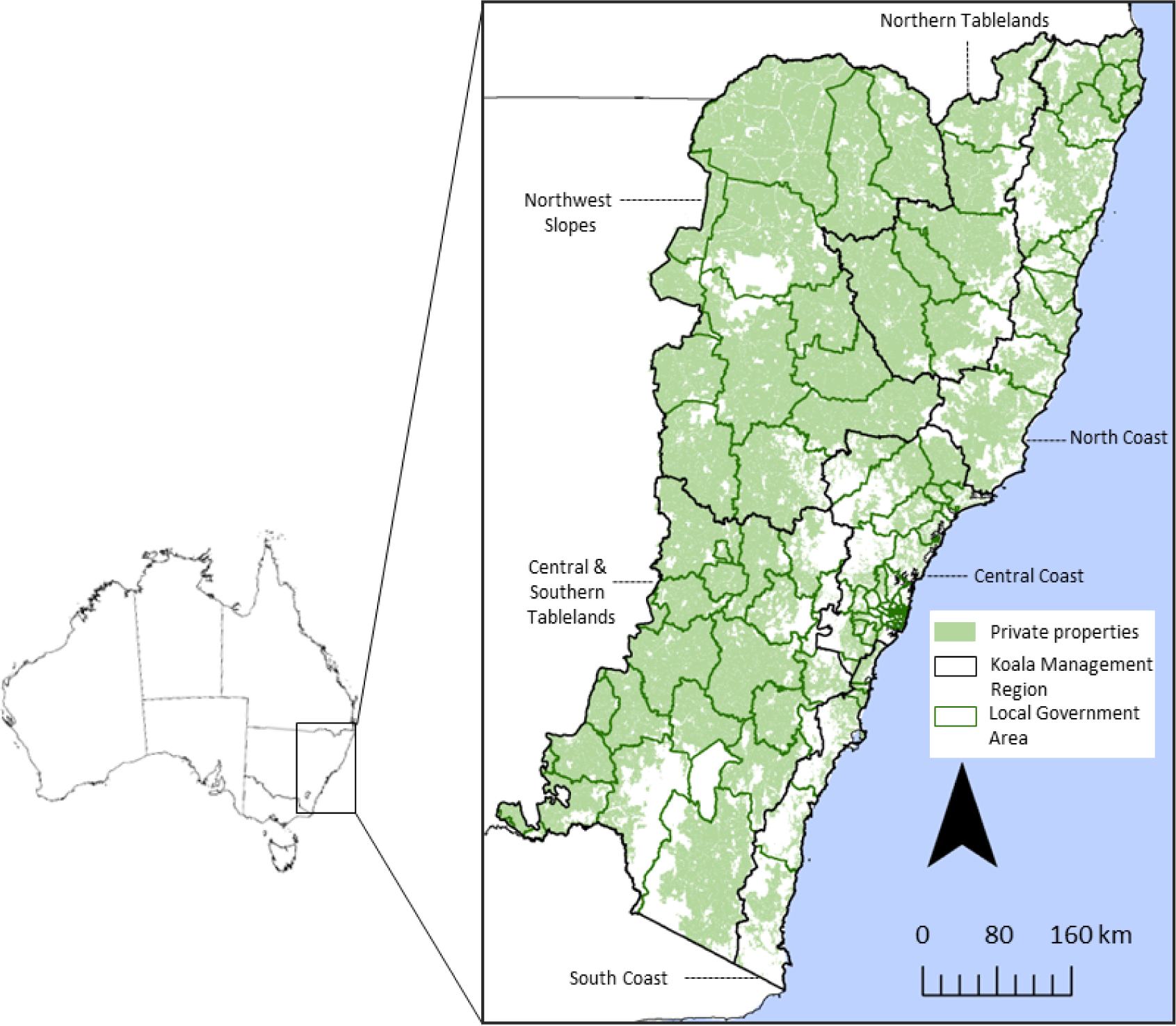
Study region encompassing the North Coast, Northern Tablelands, Northwest Slopes, Central Coast, the Central and Southern Tablelands, and the South Coast regions. Private lands (n = 199,416, that do not already have a covenant, are greater than 2 hectares in size, and were not used for intensive land-uses) are overlaid by local government areas (n = 93) (NSW Digital Cadastral Database (DCDB) 2016). See SM1.1.

### The decision problem

When BCT deploys a conservation tender it is usually highly spatially targeted to a few Local Government Areas (LGAs) rather than being delivered broadly across a large region. Hence, a key decision is where to spatially target a conservation tender to optimise delivery of key private land conservation outcomes. Once a location has been identified, there are various steps involved that BCT uses to deploy a conservation tender (see Figure 2, SM1.2). In our decision framework, we focus specifically on the conservation prioritisation problem of deciding where and how much to invest in tenders, while explicitly accounting for the tender deployment process in each location. We solve this problem assuming an LGA planning unit (the assumed scale of each tender deployment), an overall budget constraint (across all tender deployments) and an objective to maximise conservation benefit (landscape capacity to support koalas see “Koala Landscape Capacity” below - which we explore under current and future climatic conditions). In calculating the costs and conservation benefits we accounted for the likely spatial distribution and value of landholder bids in the tender process (see “Model of landholder preferences” below) the process and conservation values used by BCT to assess whether to accept bids or not (see (Biodiversity Conservation Trust 2020); and SM1.3). To achieve this we simulated the process of eliciting expressions of interest from landholders and the ranking of bids typically undertaken by BCT when deciding which bids to accept within a given tender (see SM1.2 and 1.3). For each LGA planning unit we then simulated expected conservation benefits arising from tender investment for discrete budget levels spanning the range of the approximated amounts invested in tenders previously ($1,126,260 - $32,539,448), increasing in approximately $1 million increments (see SM1.3, Figure 2 and 3). For each budget increment we ran 200 simulations (see SM1.5 for sensitivity analysis to determine number of simulations) and quantified the conservation benefits for each investment level as the average across all simulations. We assumed that a maximum of one tender could be invested in each planning unit given tenders are rarely, if ever, sequentially deployed in the same location. All monetary values were in Australian dollars ($).

**Figure 2.**
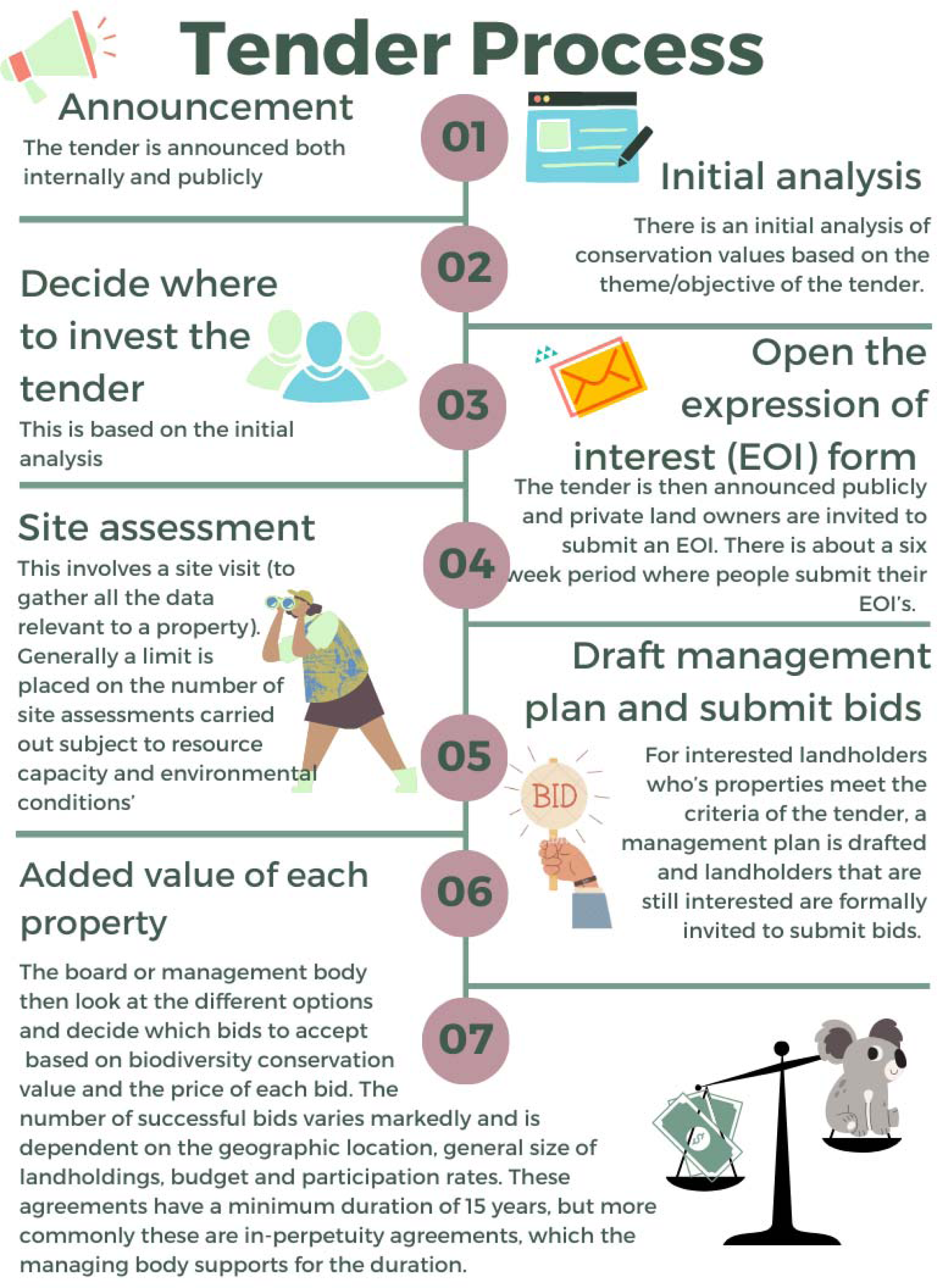
Step by step process of implementing a conservation tender. We loosely base these steps on the Biodiversity Conservation Trust’s process and note it may differ among organisations. For more details see SM1.2.

**Figure 3.**
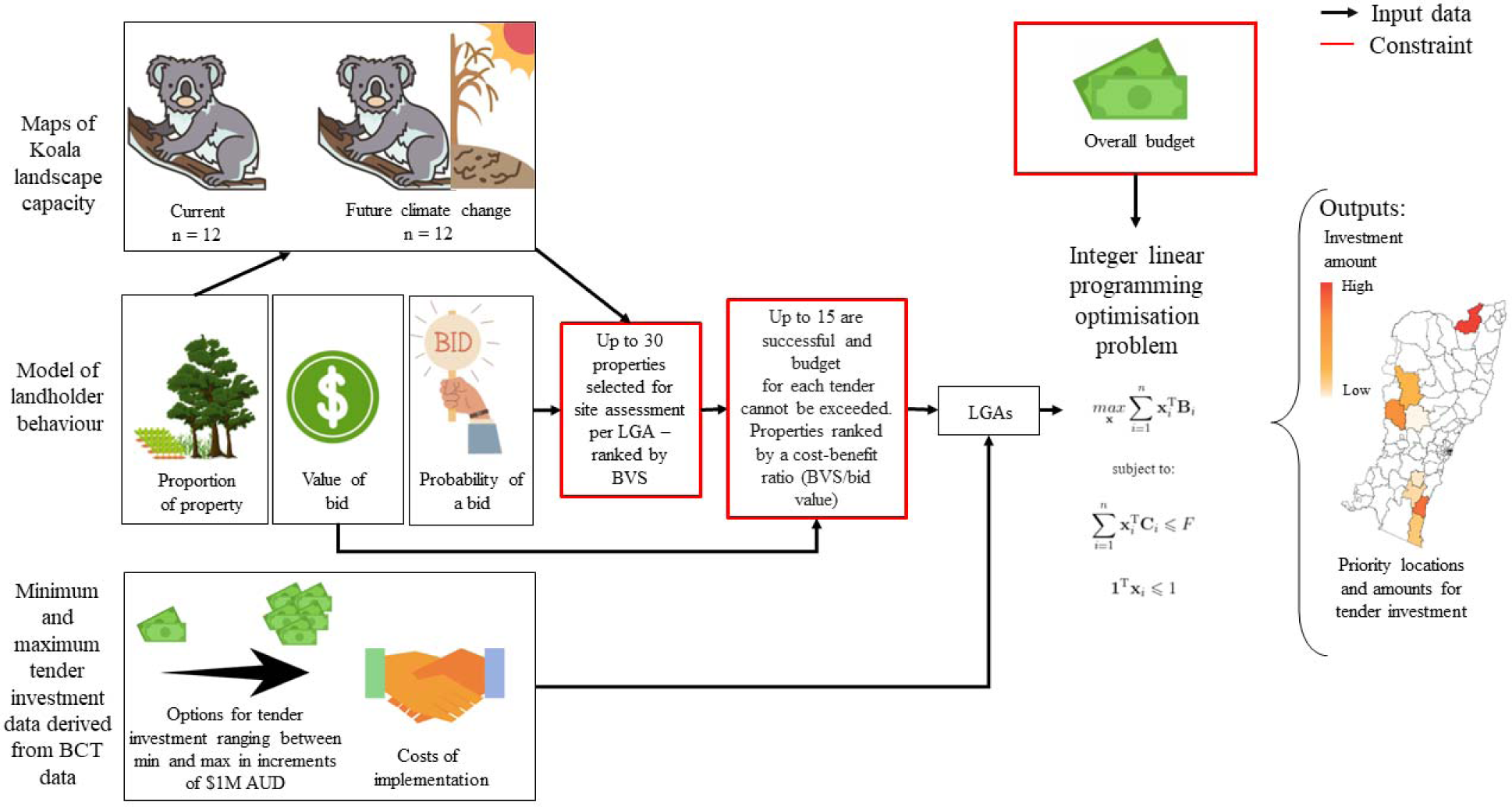
Flow diagram of the input data, process, and constraints used to develop exact spatially explicit priority solutions (outputs) by maximising the aggregated benefits of tender investments without exceeding an overall budget, solved using Gurobi Optimizer version 9.5.1 (Gurobi Optimization 2022) - for problem formulation see SM1.3. See Figure SM1.3.1 for a full flow diagram of the simulation process.

We formulated this as a mathematical optimisation problem (specifically an integer linear programming problem) and solved it using Gurobi version 9.5.1 (Gurobi Optimization 2022). See SM1.3 for mathematical formulation and more details.

### Koala Landscape Capacity

To represent conservation benefit we used koala landscape capacity under current and future climate conditions at a 90 m resolution, based on the outputs of a Rapid Evaluation of Metapopulation Persistence (REMP) model that was applied to a koala environmental niche model ((Drielsma et al. In Prep); see (Drielsma & Ferrier 2009) for methods). The REMP model outputs represent a measure of the capacity of a location to support koala populations (called “landscape capacity”) taking into account local habitat conditions, landscape connectivity, and current and future climatic conditions. For this analysis we use the latest time point before present day as the baseline (2000) and the 2070 time point to represent future climatic conditions, using the average of all 12 climate models (which represent four NARCliM v.1 General Circulation Models (CSIRO-Mk3.0, ECHAM5, MIROC3.2, and CCMA 3.1) across three Regional Climate scenarios (Evans et al. 2014)). See SM2 for more details.

### Model of landholder preferences

Based on a survey of landholders in the study region we developed empirical regression models of the proportion of their property a landholder would be willing to covenant, the probability a landholder would be willing to consider participating in a conservation covenant (which we use as a proxy for whether a landholder would place a bid in a tender), and the minimum payment per hectare per year in perpetuity a landholder would be willing to accept in return for participating in a conservation covenant on their property (which we use as proxy for the price of their bid) (see SM3 for more details). For each estimated bid value ($ ha^-1^ yr^-1^) we calculated the net present value assuming a fixed in-perpetuity annual payment using a 3.2% discount rate (the rate used in BCT’s pricing tools) (NSW Department of Planning and Environment 2023).

### Costs

In addition to the landholder payments we also accounted for the administration costs of implementing a tender. To estimate administration costs (the direct costs to BCT for each tender including staff time, staff travel and accommodation, and hours needed to develop management plans) we used the mean administrative cost of processing past tenders ($320,327.1, *SD =* $55,382.3, *n =* 11) provided by BCT. Administration costs were accounted for by subtracting the administration cost from each tender’s available budget for allocating to covenant agreements.

### Planning approaches and scenario analysis

#### Planning approaches

We considered four planning approaches to test the effect of considering landholder behaviour and the tender process in conservation planning, the approaches varied in the extent to which they considered landholder preferences and climate change. We note that landholder preferences and the tender process are closely intertwined, meaning there is dependence of our generated results on whether or not the tender process was considered. The *agnostic* approach (#1) ignores climate change, landholder preferences, and the tender process by optimising investments based only on current koala landscape capacity as conservation benefit and unimproved land value (New South Wales Government 2017)) as cost (see SM1.4). The *landholder informed* approach (#2) explicitly considers landholder preferences but ignores climate change (see SM1.3). This scenario is the same as the *agnostic* approach but incorporates landholder preferences into the tender process. The *climate informed* approach (#3) is the same as the *agnostic* approach but future koala landscape capacity under climate change is used to measure conservation benefit. The *integrated* approach (#4) combines the landholder informed and climate informed approach to consider both landholder preferences and climate change in determining optimal tender investments. To compare the performance of all approaches relative to each other we assess the priorities against the outcomes of the *integrated* approach (#4), ie. the benefit that would be realised if allocated resources were invested through a tender process in each LGA accounting for landholder preferences and future climate change. See Table 1 for summary of planning approaches.

**Table 1.**
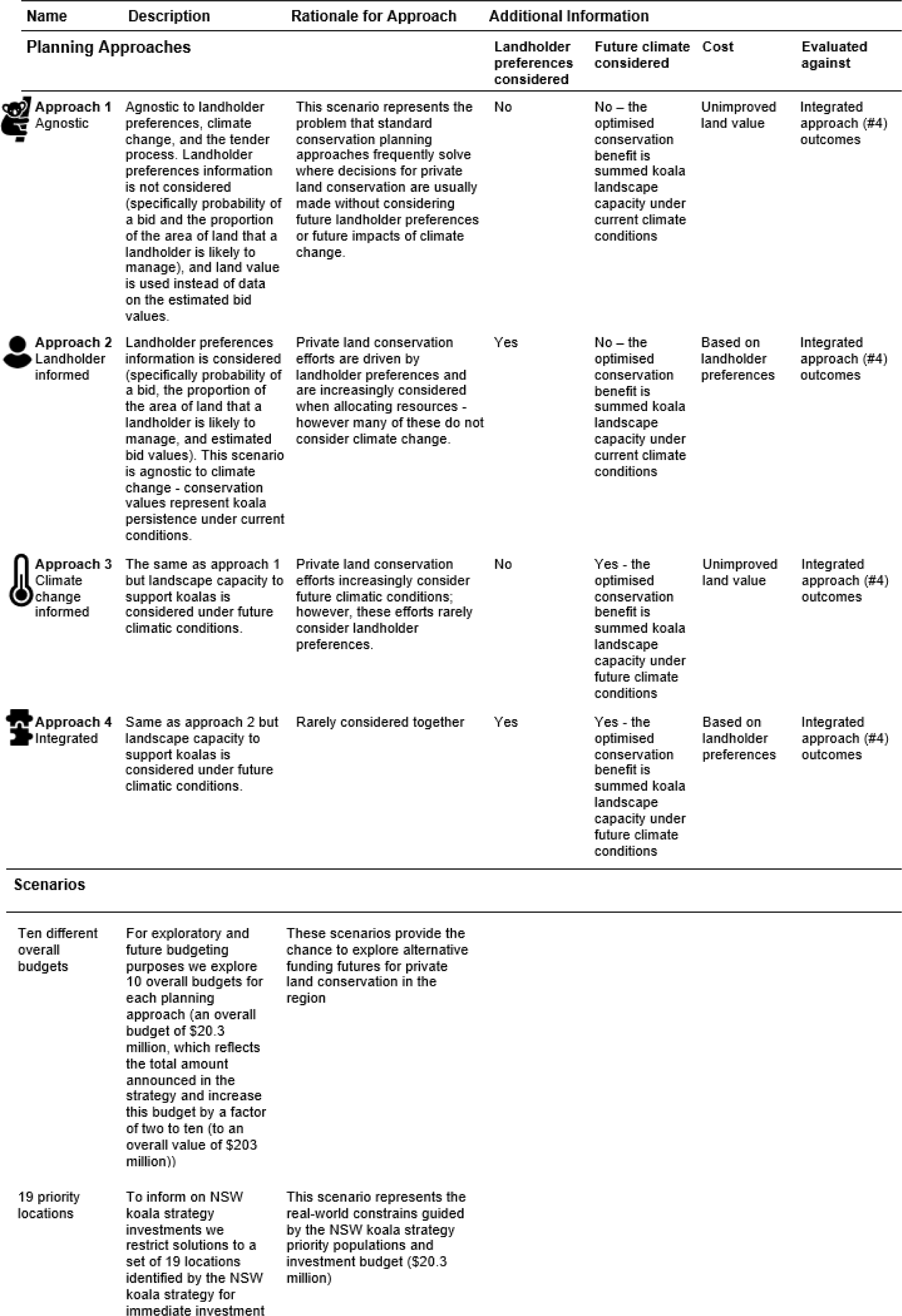
Description of the four planning evaluated approaches and decision scenarios.

| Name | Description | Rationale for Approach | Additional Information |  |  |  |
| --- | --- | --- | --- | --- | --- | --- |
| Planning Approaches |  |  | Landholder preferences considered | Future climate considered | Cost | Evaluated against |
| **Approach 1**<br>Agnostic | Agnostic to landholder preferences, climate change, and the tender process. Landholder preferences information is not considered (specifically probability of a bid and the proportion of the area of land that a landholder is likely to manage), and land value is used instead of data on the estimated bid values. | This scenario represents the problem that standard conservation planning approaches frequently solve where decisions for private land conservation are usually made without considering future landholder preferences or future impacts of climate change. | No | No – the optimised conservation benefit is summed koala landscape capacity under current climate conditions | Unimproved land value | Integrated approach (#4) outcomes |
| **Approach 2**<br>Landholder informed | Landholder preferences information is considered (specifically probability of a bid, the proportion of the area of land that a landholder is likely to manage, and estimated bid values). This scenario is agnostic to climate change - conservation values represent koala persistence under current conditions. | Private land conservation efforts are driven by landholder preferences and are increasingly considered when allocating resources - however many of these do not consider climate change. | Yes | No – the optimised conservation benefit is summed koala landscape capacity under current climate conditions | Based on landholder preferences | Integrated approach (#4) outcomes |
| **Approach 3**<br>Climate change informed | The same as approach 1 but landscape capacity to support koalas is considered under future climatic conditions. | Private land conservation efforts increasingly consider future climatic conditions; however, these efforts rarely consider landholder preferences. | No | Yes - the optimised conservation benefit is summed koala landscape capacity under future climate conditions | Unimproved land value | Integrated approach (#4) outcomes |
| **Approach 4**<br>Integrated | Same as approach 2 but landscape capacity to support koalas is considered under future climatic conditions. | Rarely considered together | Yes | Yes - the optimised conservation benefit is summed koala landscape capacity under future climate conditions | Based on landholder preferences | Integrated approach (#4) outcomes |
| Scenarios |  |  |  |  |  |  |
| Ten different overall budgets | For exploratory and future budgeting purposes we explore 10 overall budgets for each planning approach (an overall budget of \$20.3 million, which reflects the total amount announced in the strategy and increase this budget by a factor of two to ten (to an overall value of \$203 million)) | These scenarios provide the chance to explore alternative funding futures for private land conservation in the region | | | | |
| 19 priority locations | To inform on NSW koala strategy investments we restrict solutions to a set of 19 locations identified by the NSW koala strategy for immediate investment | This scenario represents the real-world constraints guided by the NSW koala strategy priority populations and investment budget (\$20.3 million) | | | | |

#### Decision Scenarios

We explore an initial decision scenario with an overall budget of $20.3 million, the total amount announced in the 2022 NSW Koala Strategy to permanently protect koala habitat over the next 5 years on private land through the BCT’s payments to private landholder programs (NSW Government 2022) (acknowledging that conservation agreements through the tender process is just one of the BCT’s mechanisms), and increase this budget by a factor of two to ten (to an overall value of $203 million). We also explore a further scenario to inform on the NSW Koala Strategy’s objectives where we used the 19 priority populations for immediate investment identified in the NSW Koala Strategy as koala populations for immediate investments (NSW Government 2022) as planning units instead of LGAs (see SM7 for more details).

## Results

### Impact of climate and landholder preferences on priority investments

We found that the agnostic approach yielded substantially different investment priorities compared to the landholder informed planning approach (Figure 4). Under the agnostic approach smaller amounts of the overall budget were allocated to more LGAs compared to the landholder informed approach where larger proportions of the budget were allocated to fewer LGAs. Optimal investments for the integrated approach were more concentrated in the LGAs surrounding Sydney, the north coast including Byron, Ballina, and Lismore, and Coffs Harbour and Nambucca. When optimising tender investment considering landholder preferences and future climatic conditions together we found priorities shift to the south-east compared to when landholder preferences are ignored (Figure 4B compared to 4D), suggesting a clear interaction between climate and landholder preference variables. This is due to the decline in koala landscape capacity to the west (SM2), restricting the cost effectiveness of payments for conservation covenants further west.

**Figure 4.**
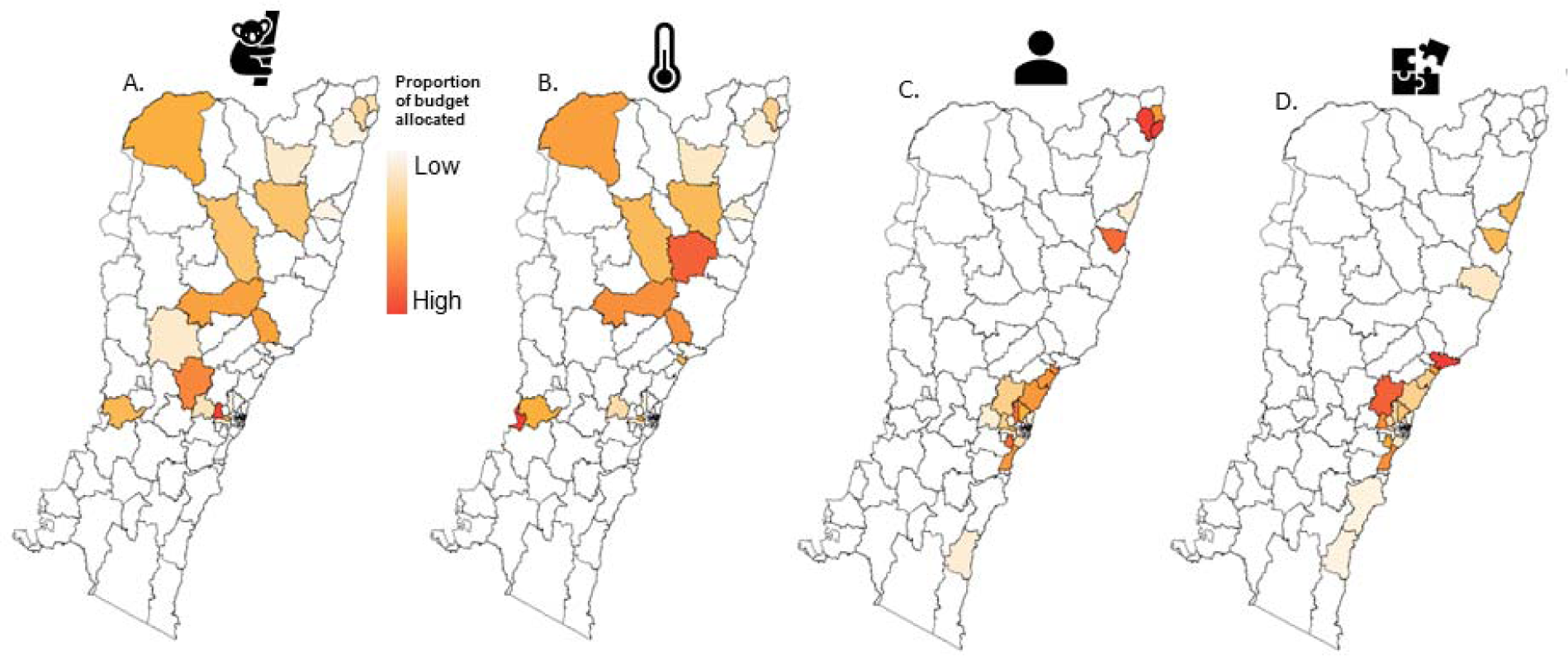
Budget allocations with an example overall budget of $101.5 million under A) the agnostic approach, B) the climate change informed approach, C) the landholder informed approach, and D) the integrated approach.

### Impact on conservation outcomes

Under every budget scenario the landholder informed and integrated approaches yielded higher conservation benefits compared to the agnostic and climate change informed approaches, on average 5.26 times higher (Figure 5, SM5). This implies that adequately accounting for landholder preferences and the tender process is far more important for achieving optimal conservation outcomes than accounting for climate change. We find that accounting for landholder preferences is so important that it obscures much of the benefit of considering climate change. When landholder preferences are considered the additional value of accounting for climate change is evident, but when they are ignored, there is little benefit to be gained (Figure 5, integrated and landholder informed approaches compared to the agnostic and climate informed approaches).

**Figure 5.**
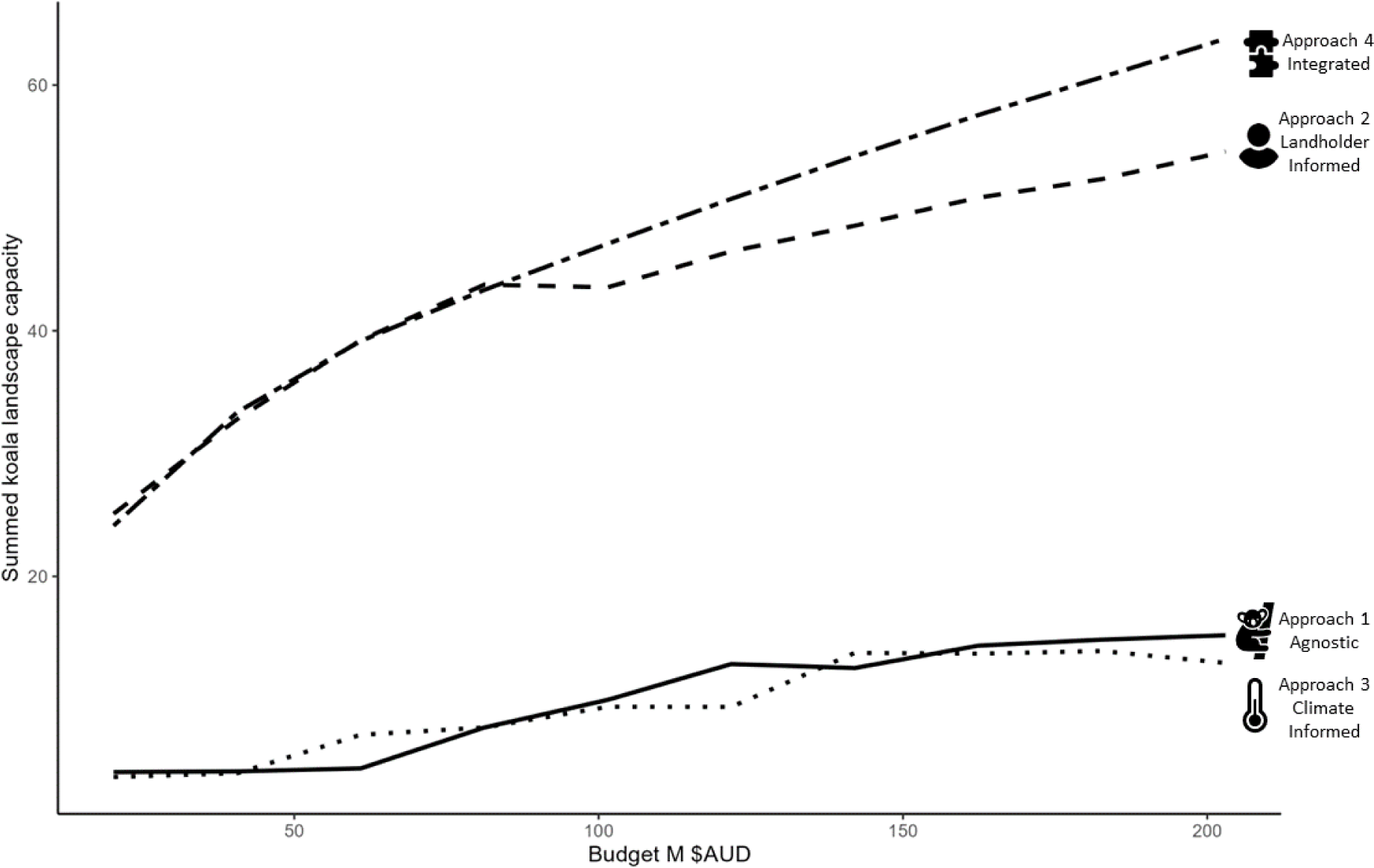
Plot showing the summed koala landscape capacity value under the agnostic approach (#1 - solid line), landholder informed approach (#2 - dashed line), climate change informed approach (#3 - dotted line), and integrated approach (#4 - two dash line). See SM5 for exact and uncertainty values.

## Discussion

Our study reveals new insights into the interaction between landholder preferences and climate change in determining conservation planning priorities and outcomes for threatened species. We found that the performance of optimal private land conservation investments for koala conservation depends more on accounting for landholder preferences and the tender process than climate change. For investment approaches that account for landholder preferences and the tender process, conservation benefits were on average 5.26 times higher than when they were ignored. Further, when landholder preferences were ignored we found no difference in benefit between approaches that considered and ignored climate change (Figure 5). This interaction means that the benefit of accounting for climate change can only be realised if landholder preferences were also accounted for; therefore, conservation planning exercises that consider climate change but not landholder preferences risk missing important conservation opportunities or even the failure of conservation actions. There is an increasing body of research dedicated to accounting for climate change in conservation planning (Jones et al. 2016), and our findings suggest that this research focus should simultaneously include the consideration of stakeholder preferences. Our findings echo support for previous studies which have called for the increased consideration of social variables throughout the planning process (Tulloch et al. 2014; Selinske et al. 2019) and for the successful implementation of planning solutions (Ban et al. 2013; Guerrero & Wilson 2017). We show this is especially important against the backdrop of climate change.

Under the climate informed approach, higher proportions of the budget are allocated towards the North Coast where koala landscape capacity is highest under future climate change (SM2) compared to the agnostic approach. However, under the integrated approach higher proportions are allocated to the Central and South Coasts. This difference is due to the fact that the assumed costs (based on unimproved land values) closer towards the North Coast are sufficiently low and ecological values sufficiently high in the climate informed approach for these areas to be prioritised, but under the integrated approach the costs in the Central and South Coasts are low enough (relatively) to switch priorities to those areas, despite the lower ecological value. This results in higher relative cost efficiency of conservation gains in the Central and South Coasts under the landholder informed approach (SM4). The implications of this are that default use of proxies for cost (for example (Strassburg et al. 2019) and (Proctor et al. 2022)) that do not account for how landholder preferences influence the true cost of conservation investments are likely to result in poor conservation outcomes (Bode et al. 2008).

Conservation tenders play a role beyond achieving conservation outcomes. In our analysis we have used information on landholder preferences to identify optimal locations for conservation tender deployment; however, this information is usually unknown or uncertain, and a conservation tender is a key mechanism for its elicitation. The information gathered can then be used to improve estimates of landholder preferences, and in the subsequent delivery of other conservation mechanisms. In this way, the BCT has leveraged its experience and information gathered to streamline its delivery of conservation programs (Biodiversity Conservation Trust 2023a). There are many conservation mechanisms (beyond those that are market-based) through which landholders might sign up for a conservation agreement such as programs that are not driven by financial incentives (for example BCT’s Conservation Partners Program’s voluntary agreements (Biodiversity Conservation Trust 2023b) and the United Kingdom’s conservation covenant agreement program (UK Government 2023)). Given our finding that accounting for the tender process is important, future decision support frameworks should focus on accounting for multiple conservation mechanisms available to a governing body, assessing the cost, benefits, and trade-offs between mechanisms, and identifying multiple priorities under each mechanism. Different mechanisms are likely to lead to quite different priorities for investment and may also interact strongly. Future work could therefore investigate how these various conservation mechanisms interact with one another, as they may either crowd each other out, enhance each other’s impact, or influence the priorities within the different mechanisms. Understanding these dynamics will be important for developing holistic and effective conservation strategies in the future.

Our analysis is potentially limited by the fact that some of the data used are proxies for data used in real tender processes. For example, to calculate values to input into the BCT’s Biodiversity Value Score calculation for each property, we used readily available metrics, such as habitat condition from the Biodiversity Indicators Program Data Package (Love et al. 2018). In reality, the Biodiversity Value Score is calculated based on intensive, and often expensive, site visits where the BCT collects data on the actual biodiversity values on a property. Additionally, the South Coast in NSW is predicted to have high koala landscape capacity in the future (SM2), but today there are few locations in this region where koalas persist (Adams-Hosking et al. 2016). Therefore, our results should be considered alongside ground-truthed information on local scale dynamics, which were not captured in our current analysis. Nonetheless, we believe the proxies used in our analysis are reasonable approximations and unlikely to influence the overall conclusions.

Our study demonstrates how making optimal decisions about where to invest in conservation tenders depends more on considering landholder preferences and tender processes than climate change. However, both are important for achieving optimal conservation outcomes as ignoring either will substantially alter tender location decisions and ultimately conservation outcomes. Our results emphasise the importance of integrating social information within decision-making frameworks for private land conservation, which not only enhances the efficacy of solutions but also aligns them with community preferences which is more likely to foster enduring positive outcomes.

## List of Supporting Material

Supporting Material 1 - Extended Methodology

Supporting Material 2 - Detail on the Koalas in the Landscape (KITL) project

Supporting Material 3 - Landholder preference models

Supporting Material 4 - Cost efficiency layers

Supporting Material 5 - Values and uncertainty for all approaches and scenarios

Supporting Material 6 - BCT rank metric and data that goes into it

Supporting Material 7 - Informing on NSW Koala Strategy Objectives

## Acknowledgements

This research was conducted with the support of funding from the Australian Research Council Linkage Project LP200100060 with financial and in-kind support from the NSW Department of Planning and Environment, the Biodiversity Conservation Trust, Tweed Shire Council, and the NSW Koala Strategy. JRR was supported by an Australian Research Council Future Fellowship (FT200100096). Figures 2 and 3 were created using https://www.canva.com/ where “All free photos, music and video files on Canva can be used for free for commercial and non-commercial use https://www.canva.com/policies/free-media-license-agreement-2022-01-03/”. The survey of landholders used to construct the landholder preference models was conducted under University of Queensland Human Ethics approval number #2019002363.

## Data availability

The annotated R code associated with these analyses is available here: https://github.com/koala-private-land/decision_support_framework. Annotated R code for landholder preference models are available here: https://github.com/koala-private-land/spatial-bid-model. These will be uploaded to a Zendo repository upon acceptance of the article. Koala REMP models which are not yet publicly available for download are available upon request from. Administrative cost data of processing past tenders cost tenders and locations of current conservation covenants are available through the Biodiversity Conservation Trust (BCT) with an appropriate licensing agreement. Survey of landholders used to construct the landholder preference models contains sensitive information and is therefore not available for use. All other data is publicly available from the source links provided in the text.

## References

Adams-Hosking, C., Grantham, H.S., Rhodes, J.R., McAlpine, C. & Moss, P.T. (2011). Modelling climate-change-induced shifts in the distribution of the koala. Wildl. Res., 38, 122–130.

Adams-Hosking, C., McAlpine, C., Rhodes, J.R., Grantham, H.S. & Moss, P.T. (2012). Modelling changes in the distribution of the critical food resources of a specialist folivore in response to climate change. Divers. Distrib., 18, 847–860.

Adams-Hosking, C., McBride, M.F., Baxter, G., Burgman, M., de Villiers, D., Kavanagh, R., Lawler, I., Lunney, D., Melzer, A., Menkhorst, P., Molsher, R., Moore, B.D., Phalen, D., Rhodes, J.R., Todd, C., Whisson, D. & McAlpine, C.A. (2016). Use of expert knowledge to elicit population trends for the koala (Phascolarctos cinereus). Divers. Distrib., 22, 249–262.

Ban, N.C., Mills, M., Tam, J., Hicks, C.C., Klain, S., Stoeckl, N., Bottrill, M.C., Levine, J., Pressey, R.L., Satterfield, T. & Chan, K.M.A. (2013). A social–ecological approach to conservation planning: embedding social considerations. Front. Ecol. Environ., 11, 194–202.

Biodiversity Conservation Trust. (2018). Conservation Tenders - Guide for Landholders. https://www.bct.nsw.gov.au/sites/default/files/2018-05/Conservation_Tenders_Guide_for_Landholders_updated.pdf

Biodiversity Conservation Trust. (2020). Biodiversity Conservation Trust Assessment Metric. https://www.bct.nsw.gov.au/sites/default/files/2020-09/BCT%20Assessment%20Metric%20Web%20Version%20September%202020.pdf

Biodiversity Conservation Trust. (2023a). Investing in Private Land Conservation. NSW Government. https://www.bct.nsw.gov.au/sites/default/files/2023-07/conservation-management-program-2023-2027.pdf

Biodiversity Conservation Trust. (2023b). Conservation Partners Program. https://www.bct.nsw.gov.au/conservation-partners-program

Blanch, S., Sweeney, O. & Pugh, D. (2022). The NSW Koala Strategy: ineffective, inadequate and expensive. An assessment of the NSW Koala Strategy against recommendations made in the Report of the Independent Review into the Decline of Koala Populations in Key Areas of NSW. https://npansw.org.au/wp-content/uploads/2019/03/koala-strategy-update.pdf

Bode, M., Wilson, K.A., Brooks, T.M., Turner, W.R., Mittermeier, R.A., McBride, M.F., Underwood, E.C. & Possingham, H.P. (2008). Cost-effective global conservation spending is robust to taxonomic group. Proc. Natl. Acad. Sci. U. S. A., 105, 6498–6501.

Cortés Capano, G., Toivonen, T., Soutullo, A. & Di Minin, E. (2019). The emergence of private land conservation in scientific literature: A review. Biol. Conserv., 237, 191–199.

Department of the Environment. (2022). Phascolarctos cinereus (combined populations of Qld, NSW and the ACT) — Koala (combined populations of Queensland, New South Wales and the Australian Capital Territory). http://www.environment.gov.au/cgi-bin/sprat/public/publicspecies.pl?taxon_id=85104

Drielsma, M. & Ferrier, S. (2009). Rapid evaluation of metapopulation persistence in highly variegated landscapes. Biol. Conserv., 142, 529–540.

Drielsma, M.J., Love, J., Thapa, R., Thonell, J. & Beaumont, L.J. (In prep). Koalas in the landscape (KITL). Landscape capacity to support Koala populations through climate change. Department of Planning and Environment, NSW Government.

Fitzsimons, J.A. & Carr, C.B. (2014). Conservation covenants on private land: issues with measuring and achieving biodiversity outcomes in Australia. Environ. Manage., 54, 606–616.

Fitzsimons, J. & Cooke, B. (2021). Key questions for conservation tenders as a means for delivering biodiversity benefits on private land. Ecol. Manage. Restor., 22, 110–114.

Gooden, J. & ’t Sas-Rolfes, M. (2020). A review of critical perspectives on private land conservation in academic literature. Ambio, 49, 1019–1034.

Government of New South Wales. (2016). Biodiversity Conservation Act 2016 No 63. https://legislation.nsw.gov.au/view/html/inforce/current/act-2016-063

Graham, E.M., Reside, A.E., Atkinson, I., Baird, D., Hodgson, L., James, C.S. & VanDerWal, J.J. (2019). Climate change and biodiversity in Australia: a systematic modelling approach to nationwide species distributions. Australasian Journal of Environmental Management, 26, 112–123.

Guerrero, A.M. & Wilson, K.A. (2017). Using a social-ecological framework to inform the implementation of conservation plans. Conserv. Biol., 31, 290–301.

Gurobi Optimization, L.L.C. (2022). Gurobi optimizer reference manual.

IPCC. (2023). Climate Change 2023 Synthesis Report Summary for Policymakers. In: Climate Change 2023: Synthesis Report. A Report of the Intergovernmental Panel on Climate Change. Contribution of Working Groups I, II and III to the Sixth Assessment Report of the Intergovernmental Panel on Climate Change [Core Writing Team, H. Lee and J. Romero (eds.)]. IPCC, Geneva, Switzerland, 36 pages.

Jones, K.R., Watson, J.E.M., Possingham, H.P. & Klein, C.J. (2016). Incorporating climate change into spatial conservation prioritisation: A review. Biol. Conserv., 194, 121–130.

Kamal, S., Grodzińska-Jurczak, M. & Brown, G. (2015). Conservation on private land: a review of global strategies with a proposed classification system. J. Environ. Planning Manage., 58, 576–597.

Kearney, S.G., Carwardine, J., Reside, A.E., Adams, V.M., Nelson, R., Coggan, A., Spindler, R. & Watson, J.E.M. (2022). Saving species beyond the protected area fence: Threats must be managed across multiple land tenure types to secure Australia’s endangered species. Conservat Sci and Prac, 4.

Love, J., Drielsma, M.J., Williams, K. & Thapa, R. (2018). Data package for habitat condition indicators; 3.1a ecological condition, 3.1b ecological connectivity and 3.1c ecological carrying capacity. NSW Office of Environment and Heritage.

Maxwell, S.L., Cazalis, V., Dudley, N., Hoffmann, M., Rodrigues, A.S.L., Stolton, S., Visconti, P., Woodley, S., Kingston, N., Lewis, E., Maron, M., Strassburg, B.B.N., Wenger, A., Jonas, H.D., Venter, O. & Watson, J.E.M. (2020). Area-based conservation in the twenty-first century. Nature, 586, 217–227.

New South Wales Government. (2017). Bulk land value information available on the Valuation Services Portal.

Nielsen, A.S.E., Strange, N., Bruun, H.H. & Jacobsen, J.B. (2017). Effects of preference heterogeneity among landowners on spatial conservation prioritization. Conserv. Biol., 31, 675–685.

NSW Department of Planning and Environment. (2023). Total Fund Deposit and discount rate. https://www.environment.nsw.gov.au/topics/animals-and-plants/biodiversity-offsets-scheme/generate-credits-biodiversity-stewardship-agreement/total-fund-deposit#:~:text=The%20most%20recent%20review%20was%20undertaken%20in%20July,Stewardship%20Agreements%20entered%20into%20from%2015%20August%202022.

NSW Digital Cadastral Database (DCDB). (2016). NSW Administrative Boundaries. NSW Administrative Boundaries. https://datasets.seed.nsw.gov.au/dataset/nsw-administrative-boundaries

NSW Government. (2022). NSW Koala Strategy - Towards doubling the number of koalas in New South Wales by 2050. https://www.environment.nsw.gov.au/-/media/OEH/Corporate-Site/Documents/Animals-and-plants/Threatened-species/koala-strategy-2022-220075.pdf

Proctor, C.A., Schuster, R., Buxton, R.T. & Bennett, J.R. (2022). Prioritization of public and private land to protect species at risk habitat. Conserv. Sci. Pract., 4.

Rolfe, J., Schilizzi, S. & Iftekhar, M.S. (2022). Increasing environmental outcomes with conservation tenders: The participation challenge. Conservation Letters, 15, e12856.

Selinske, M.J., Howard, N., Fitzsimons, J.A., Hardy, M.J. & Knight, A.T. (2022). “Splitting the bill” for conservation: Perceptions and uptake of financial incentives by landholders managing privately protected areas. Conserv. Sci. Pract., 4, e12660.

Selinske, M.J., Howard, N., Fitzsimons, J.A., Hardy, M.J., Smillie, K., Forbes, J., Tymms, K. & Knight, A.T. (2019). Monitoring and evaluating the social and psychological dimensions that contribute to privately protected area program effectiveness. Biol. Conserv., 229, 170–178.

Strassburg, B.B.N., Beyer, H.L., Crouzeilles, R., Iribarrem, A., Barros, F., de Siqueira, M.F., Sánchez-Tapia, A., Balmford, A., Sansevero, J.B.B., Brancalion, P.H.S., Broadbent, E.N., Chazdon, R.L., Filho, A.O., Gardner, T.A., Gordon, A., Latawiec, A., Loyola, R., Metzger, J.P., Mills, M., Possingham, H.P., Rodrigues, R.R., Scaramuzza, C.A. de M., Scarano, F.R., Tambosi, L. & Uriarte, M. (2019). Strategic approaches to restoring ecosystems can triple conservation gains and halve costs. Nat. Ecol. Evol., 3, 62–70.

Tulloch, A.I.T., Tulloch, V.J.D., Evans, M.C. & Mills, M. (2014). The value of using feasibility models in systematic conservation planning to predict landholder management uptake. Conserv. Biol., 28, 1462–1473.

UK Government. (2023). Getting and using a conservation covenant agreement. https://www.gov.uk/guidance/getting-and-using-a-conservation-covenant-agreement

Whitten, Wünscher & Shogren. (2017). Conservation tenders in developed and developing countries− status quo, challenges and prospects. Land use policy, 63, 552–560.

