## Supporting Material for "Optimal investments in private land conservation depend more on landholder preferences than climate change"

#### Supporting Material 1 - Extended Methodology

##### 1.1 Delineating property boundaries

We used the NSW Property Theme Dataset (NSW Government 2022a) to delineate private property boundaries.

We used the lots spatial unit from the 2019 NSW Digital Cadastre Dataset to characterise parcels of land created on a survey plan and the properties spatial unit from the 2022 NSW Property Theme Dataset to characterise areas of land under common ownership (noting that properties can consist of multiple lots). We first excluded all lots on crown land, all state forests, and all protected areas in the 2018 Collaborative Australian Protected Areas Database (CAPAD) (except for "private nature reserves''). We then excluded all lots with an area under 2 ha (a threshold below which we considered would be too small to be of interest for private land conservation activities as this reflects the smallest agreement BCT has entered as part of a koala tender historically), areas identified as roads, road corridors, and rail corridors, lots under intensive land uses (assuming there is little chance these would be targeted for private land conservation), and lots covered by water (assuming little benefit for koalas).

Specifically, excluded intensive land uses (based on the most common land use within each lot) were:

Manufacturing and Industrial (5.3.0)

Urban Residential (5.4.1)

Services (5.5.0)

Utilities (5.6.0)

Transport and Communication (5.7.0)

Mining (5.8.0)

Waste Treatment and Disposal (5.9.0)

Excluded water land uses (based on the most common land use within each lot) were:

Lake (6.1.0)

Reservoir/dam (6.2.0)

River (6.3.0)

Channel/aqueduct (6.4.0)

Estuary/coastal waters (6.6.0)

Land uses were taken from the NSW Landuse 2017 v1.2 spatial data set (<https://datasets.seed.nsw.gov.au/dataset/nsw-landuse-2017-v1p2-f0ed>) and land use codes are based on secondary or tertiary Australian Land Use and Management Classification Version 8 categories (<https://www.agriculture.gov.au/abares/aclump/land-use/alum-classification>).

Finally, each lot was assigned to a common property based on the centroid of the lot and then dissolved by property to define spatial representations of properties used in the analysis.

##### 1.2 Step by step process of implementing a conservation tender

Here we outline the broad step by step process of implementing a conservation tender that we aimed to simulate within the prioritisation process. We loosely base these steps on the Biodiversity Conservation Trust’s process and note it will differ among organisations.

1. **Announcement** – The tender is announced both internally and publicly.
2. **Initial analysis** – There is an initial analysis of conservation values of different areas based on the theme/objective of the tender.
3. **Decide where to invest in and deploy the conservation tender** – This is based on the initial analysis.
4. **Open the expression of interest (EOI) form** – The location of the tender is then announced and private land owners are invited to submit an EOI. There is about a six week period where people submit their EOI’s.
5. **Site assessment of the properties who submitted EOI’s** – This involves a site visit (to gather all the data relevant to a property). Generally a limit is placed on the number of site assessments carried out subject to resource capacity and environmental conditions. It is important to manage the expectations of successful bids.
6. **Draft management plan and submit bids** – For interested landholders who’s properties meet the criteria of the tender, a management plan is drafted and landholders that are still interested are formally invited to submit bids.
7. **The added value of each property to the agreement/protected area portfolio** – There is another biodiversity valuation of each property to help decide which bid to accept. In the case of the Biodiversity Conservation Trust this is done using the Biodiversity Valuation Score (BVS). The BVS is an assessment metric which synthesises information on conservation value, duration, risk, and area to determine best value for money sites in BCT’s conservation management programs. For more details see (Biodiversity Conservation Trust 2020). The BVS is then used to determine the Biodiversity Value Index (BVI), or the cost-benefit ratio, for each property. The board or management body then looks at the different options and decides which bids to accept based on biodiversity conservation value and the price of each bid. The number of successful bids varies markedly and is dependent on the geographic location, general size of landholdings, budget and participation rates. These agreements have a minimum duration of 15 years, but more commonly these are in-perpetuity agreements, which the managing body supports for the duration.

##### 1.3 Problem formulation for the landholder informed and integrated approaches

In the initial tender planning stages (ie. deciding where to deploy a conservation tender) the likely outcomes of later stages should be considered which are influenced by landholder preferences (e.g., which landowners are likely to bid and how much), which bids will likely be accepted (considering the conservation benefits of each property now and into the future), and what the likely costs are going to be. Systematic conservation planning (a structured systematic approach to conservation) is one mechanism to inform decisions about which regions to deploy market-based mechanisms (Hajkowicz et al. 2007, 2008; Marinoni et al. 2008, 2009). Decision support frameworks exist for deciding which bids to accept as part of a tender (Hajkowicz et al. 2007; Marinoni et al. 2009); however, to optimally identify and make informed choices about which region to deploy a conservation tender in it is necessary to consider the expected costs and benefits of the entire process compared across space considering the above factors. In particular, the bids that are accepted in the final stages of the decision-making processes ultimately drive cost effectiveness and feasibility and therefore must be considered in the initial decision stages.

We frame the decision problem as how much to optimally invest in a tender in each LGA assuming an overall budget constraint to maximise conservation benefit (highest koala landscape capacity value) while accounting for the spatial distribution of bids, which depends on landholder preferences, and the spatial distribution of conservation values (which depend on current and future ecological states) at the property level within planning units. We assume that a maximum of one tender can be invested in each planning unit given tenders are rarely, if ever, implemented simultaneously in the same location. To account for the effect of the spatial distributions of bids, conservation values, and conservation outcomes at the property level we integrate a model of landholder preferences with an ecological model of koala landscape capacity now and across a range of future climate projections. We explicitly embed a simulation of the ranking process typically undertaken by BCT when deciding which properties to select within a given tender (see SM1.2). At the individual tender level we simulate expressions of interest from landholders, the selection of a subset of these properties for a site assessment and invitation to submit a tender bid, the ranking of these bids based on cost efficiency, and selecting the highest ranked until either the maximum number of properties that can be accepted is reached or the budget available is reached. We simulate tender benefits for a range of budgets spanning the lowest approximate amount ($1,126,260) that had been previously invested in a tender by BCT, increasing in approximately $1 million increments to the highest approximate ($32,539,448) previous investment value for a single tender by BCT. We formulate this as an integer linear programming optimisation problem as follows:

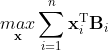
 (1)

subject to:

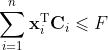

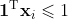

where the binary decision variable **x***_i_* is a vector (of length equal to the number of alternative budget levels) representing the budget level invested in for planning unit *i* (with an entry of 1 for the budget level chosen and 0 otherwise), **B***_i_* is a vector representing the conservation benefit of investing at each budget level in planning unit *i*, **C***_i_* is a vector representing the economic cost of investing at each budget level in planning unit *i*, *F* is the overall economic budget available, **1** is a vector of ones of length equal to the number alternative budget levels, and *n* is the number of planning units (in this case planning units are LGAs). T signifies the transpose of a matrix. The first constraint ensures the total economic budget is not exceeded and the second constraint ensures that only one budget level is selected for each planning unit.

First, we simulate the conservation benefit derived from the tender process (for each planning unit *i* and budget increment *d*) through the following steps. First, to simulate the landholders that would put in expressions of interest we selected properties based on a Bernoulli distribution with probability equal to the expected probability that a landholder in each property would consider a covenant derived from the landholder preference models (see SM3 for more details). We assume that all landholders who submit an expression of interest place a bid if invited to do so through the tender simulation process below (Figure S1.3.1).

Second, expressions of interest were then ranked according to BCT’s Biodiversity Value Score (BVS (Biodiversity Conservation Trust 2020)), with the highest 30 assumed to be selected for a site assessment. BVS is calculated as:

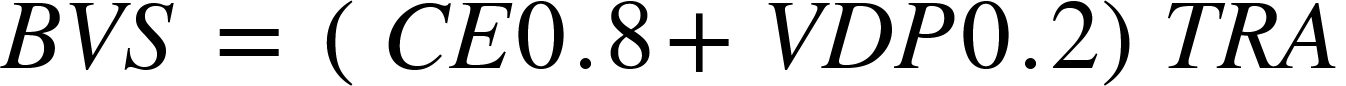
 (2)

where *C* is ecological condition based on a property’s current and future predicted condition based on management actions agreed by the landholder to be undertaken at a site, and *E* is the status based on a summed scoring system of species occurring within the property and their respective conservation status. In lieu of conducting a comprehensive site assessments for each property we used the mean Condition from the Biodiversity Indicators Program Data Package (Love et al. 2018), and given the koala-focus of the tender we estimate *E* for each property based on a the koala landscape capacity values of each property (see SM2). The variable *V* is the landscape context representing how well-connected the property is to surrounding habitat (defined as the mean Carrying Capacity from the Biodiversity Indicators Program Data Package (Love et al. 2018)) within a 1500m buffer, *D* is proximity to other properties under a conservation covenant according to BCT’s Site Assessment Geodatabase (allocated a score between 1-100), and *P* is proximity to protected areas (NSW Government 2022b) allocated a score between 1-100 (NSW Government 2022b). *T* represents the duration of the agreement with in-perpetuity agreements receiving a value of 1 (which we assume for all properties), *R* represents risk of habitat conversion derived from land and soil capability as a proxy (DPE 2021) and based on a scoring system (a class of ≥3 = 6, 4=3, 5-6 = 2, and 7-8 = 1), and *A* is the area of the property. In actual tender settings a site assessment is conducted which involves a site visit (to gather data relevant to a property), and generally there is a limit on how many site assessments can be carried out – which we assume is limited to 30. Therefore, we take the 30 properties with the highest BVS (if there are >30) to the next step.

Third, a management plan is drafted, which property owners can review and decide on whether placing a bid is still within their interest. We assume that all properties selected in the previous step are assumed to make a bid and the cost efficiency is calculated. The expected monetary value that each landholder is likely to bid (*p_ij_*) was obtained from the landholder preference models (SM3) and then used to determine the cost efficiency of investing in each property:

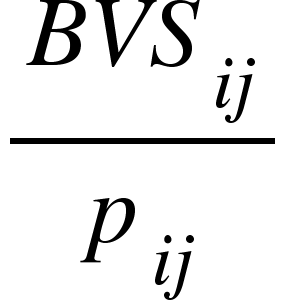
(3)

Finally, properties were ranked by this cost-benefit ratio to identify those that would likely be selected for each planning unit *i*, until either 15 properties are selected or the maximum budget available for each planning unit *i* under each budget increment *d* is reached in each simulation *k*. See Figure S1.3.1 for flow diagram of this simulation and how it aligns with input data and the tender process.

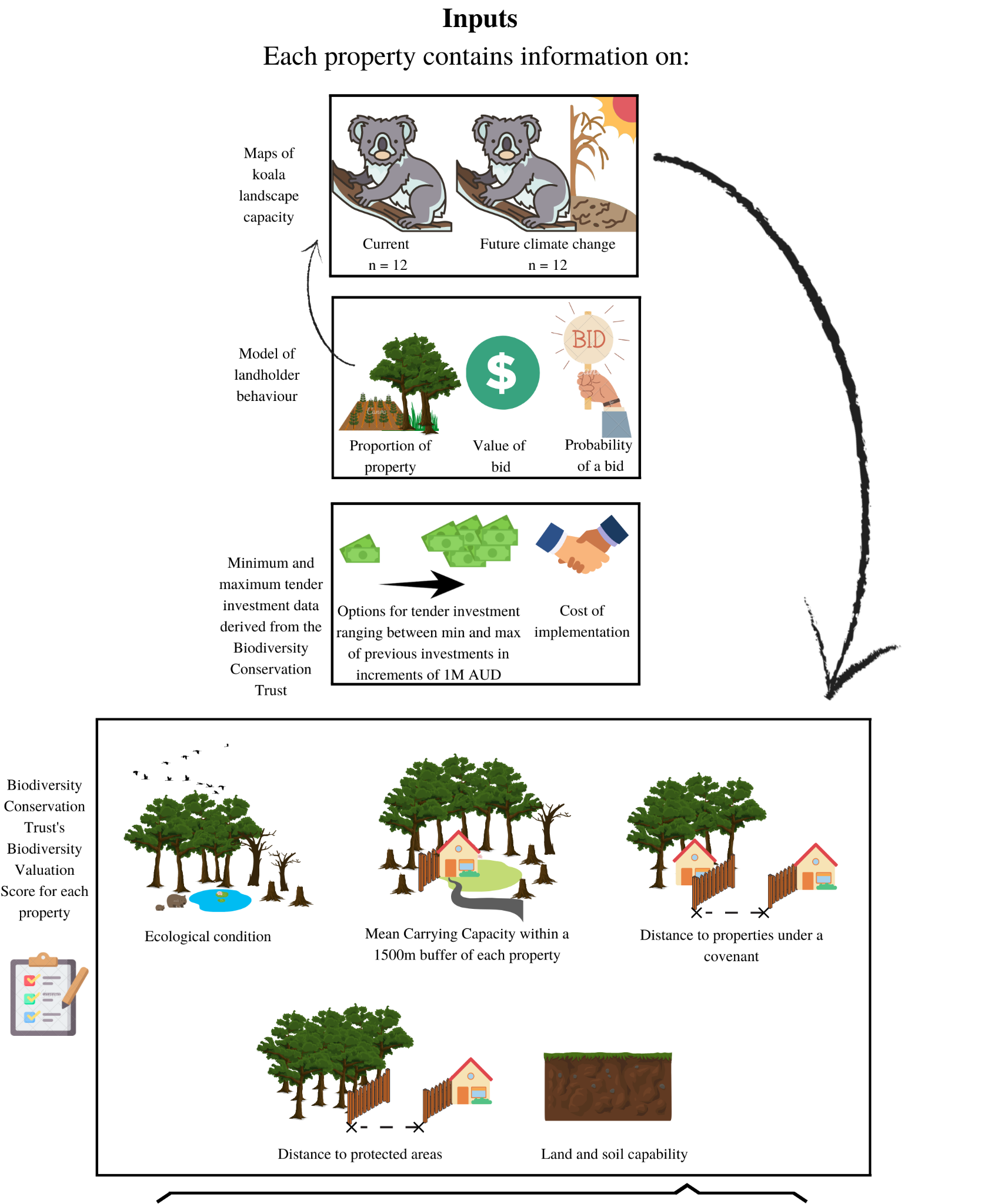

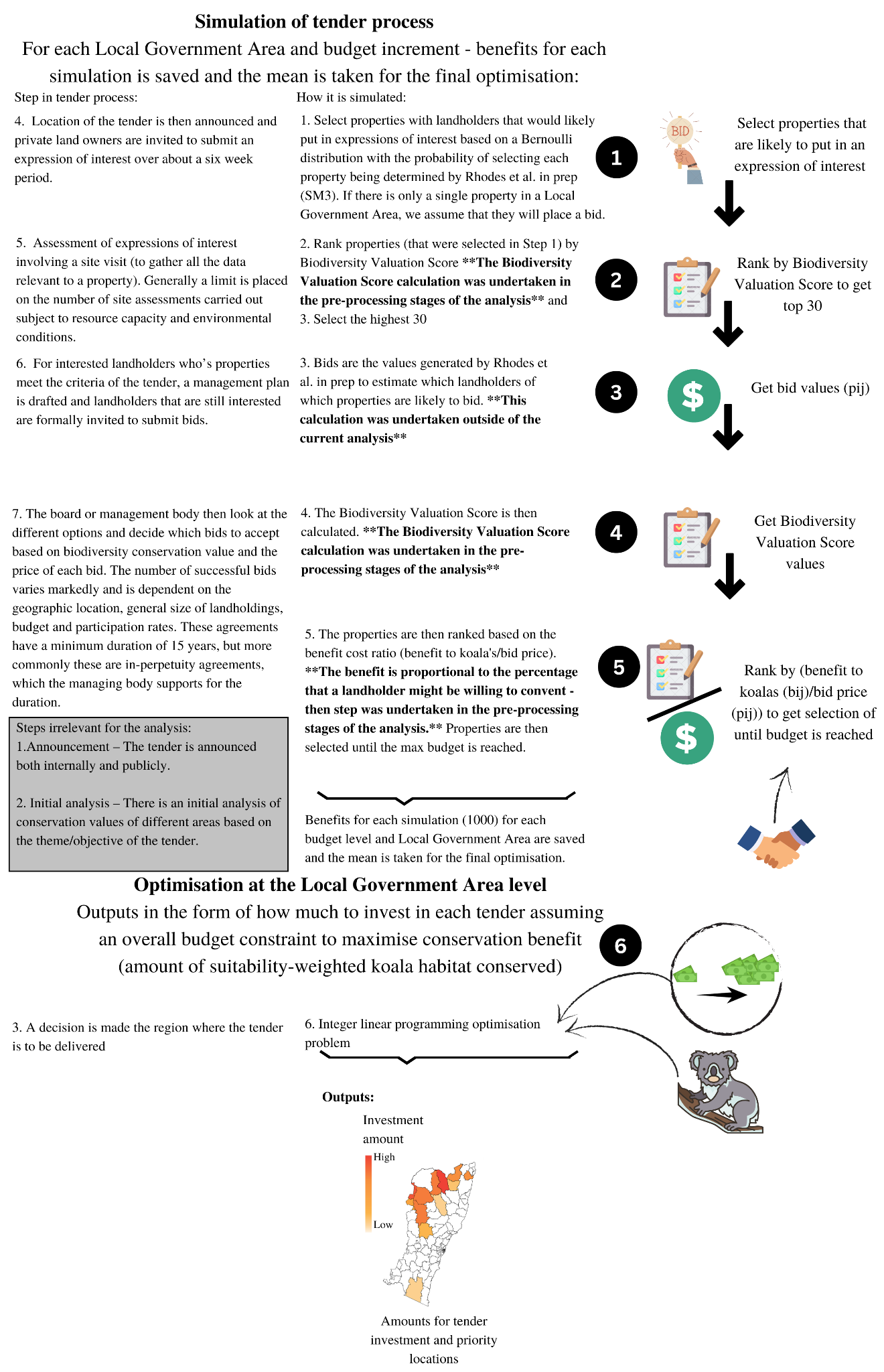

**Figure S1.3.1** - Figure showing how the simulation used in this analysis aligns with the tender process

The expected benefits in equation 1 **B***_i_*, depends on the properties for which bids are accepted in the planning unit that the tender is deployed in and the gain in ecological values that arises from each new covenant (determined through the tender simulation process). The expected benefits for a tender in a planning unit for a given budget level were calculated as

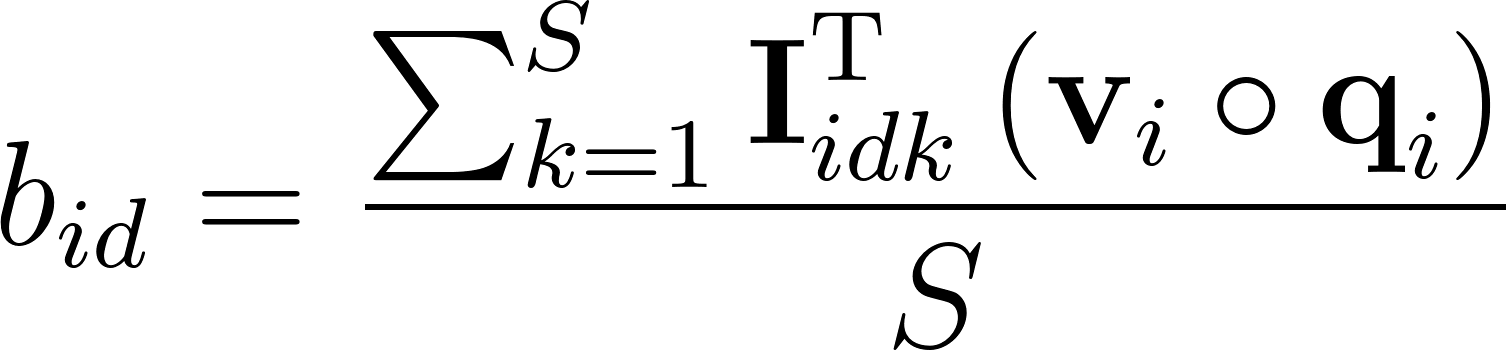
<https://www.codecogs.com/eqnedit.php?latex=b_%7Bid%7D%3D%5Cfrac%7B%5Csum_%7Bk%3D1%7D%5E%7BS%7D%5Ctextbf%7BI%7D_%7Bidk%7D%5E%7B%5Ctextrm%7BT%7D%7D%5Cleft%20(%5Ctextbf%7Bv%7D_%7Bi%7D%20%5Ccirc%20%5Ctextbf%7Bq%7D_%7Bi%7D%5Cright%20)%7D%7BS%7D#0>	(4)

where *b_id_* is the expected conservation benefit for planning unit *i* for budget increment *d* (i.e., entry *d* in the vector **B***_i_* in equation 1), **I***_idk_* is a vector (of length equal to the number of properties in planning unit *i*) containing binary indicators of which properties were selected in planning unit *i* for budget level *d* and simulation *k*, **v***_i_* is a vector (of length equal to the number of properties in planning unit *i*) containing the summed koala landscape capacity in each property, **q***_i_* is a vector (of length equal to the number of properties in planning unit *i*) containing the expected proportion of each property that a landholder would willing to covenant obtained from the landholder preference models (SM3), and *S* is the number of simulations. T signifies the transpose of a matrix and ○ signifies component-wise matrix multiplication. Using equation 4 we simulated *b_id_* for each budget increment and each LGA planning unit 200 times chosen based on the point of diminishing returns - see SM1.5.

Koala landscape capacity values (**v***_i_*) were represented by baseline (2000) climatic conditions under the landholder informed approach, and 2070 conditions under the integrated management approach. Values are the mean of 12 climate scenarios (four General Circulation Models (CSIRO-Mk3.0, ECHAM5, MIROC3.2, and CCMA 3.1) under three Regional Climate scenarios). See SM2 for a more detailed description of the models.

##### 1.4 Problem formulation under the agnostic and climate change informed approaches

Under the agnostic and climate change informed approaches we make the following adjustments to the problem formulation. We chose the selected properties in equation 4 (**I***_idk_*) based instead on an LGA level standard prioritisation with unimproved land value for each property (New South Wales Government 2017) as cost (*c*) and a budget constraint (each budget increment *d*). To achieve this we solved the following decision problem

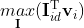
(5)

subject to:

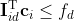

where **I***_id_* is the vector (of length equal to the number of properties in planning unit *i*) containing binary indicators of which properties are selected in planning unit *i* for budget level *d*, **v***_i_* is a vector (of length equal to the number of properties in planning unit *i*) containing the summed koala landscape capacity in each property, **c**_i_ is a vector (of length equal to the number of properties in planning unit *i*) of the unimproved land values for planning unit *i*, and *f_d_* is the budget level that corresponds to budget increment *d*. Note that there is no need for *k* (the simulation index) as in equation 4 as *S* = 1 (i.e., which properties are selected is deterministic and therefore there is no need for multiple simulations) and we also set the expected proportion of properties covenanted to 1 (i.e., setting all entries in the vector **q***_i_* to 1 from equation 4 and assuming entire properties are protected if selected). The constraint ensures the sum of the costs do not exceed budget level. This optimisation was run for each planning unit and for each budget increment and then used to calculate **B**_i_ and **C**_i_ in equation 1 accordingly and then the program level optimisation was run as for the landholder informed and integrated approaches. To obtain comparable conservation benefits (what you would actually get if the optimal investment in each planning unit was invested in conservation tenders) we ran the allocated budgets through the simulation process described in 1.3.

Again, koala landscape capacity values (**v***_i_*) were represented by baseline (2000) climatic conditions under the agnostic approach, and 2070 conditions under the climate informed approach. Values are the mean of 12 climate scenarios (four General Circulation Models (CSIRO-Mk3.0, ECHAM5, MIROC3.2, and CCMA 3.1) under three Regional Climate scenarios). See SM2 for a more detailed description of the models.

##### 1.5 Sensitivity analysis to identify the sufficient number of simulations

To determine an appropriate number of simulations to estimate the benefit functions (equation 2), we plotted the mean conservation benefit (ie. summed koala landscape capacity; across all tender budget increments) standard error against the number of simulations used, using an overall budget of $20.3 million AUD. Based on this we decided to use 200 simulations throughout as this is roughly the point at which little gain in decreased standard error is achieved by increasing the number of simulations used (Figure S1.5.1).

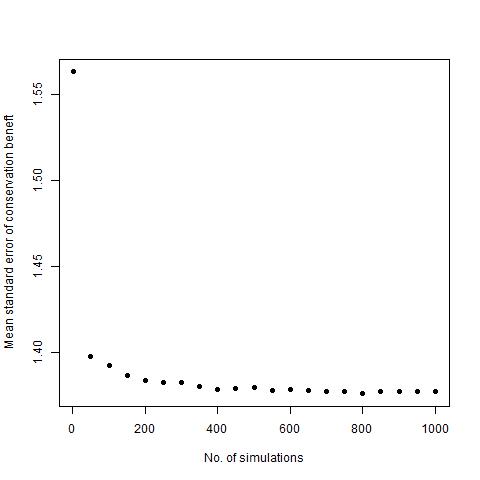

**Figure S1.5.1.** Mean (across all tender budget increments) standard error of conservation benefit (koala landscape capacity) for increasing numbers of simulations

#### Supporting Material 2 - Koalas in the Landscape (KITL) project

The Koala Landscape Capacity dataset represents the outputs of a Rapid Evaluation of Metapopulation Persistence (REMP) model ({Drielsma et al. n.d.) that was developed to assess the capacity of NSW landscapes to support koala populations. It integrates the suitability of koalas habitat across NSW and its spatial configuration, in the contemporary epoch as well as considering the likely effects of climate change up to 2070. Output data both evaluates future landscape capacity and provides guidance to conservation action. The koala landscape capacity model is developed over three main steps (Figure S2.1):

1. the development of three sub-models (the Koala Bio-Climatic Suitability Model (KBSM), The Koala Tree Species Model (KTM), and Woody Extent (WE))
2. the combining of the sub-models to derive of the Koala Environmental Niche Model (KENM)
3. applying the REMP model to the KENM

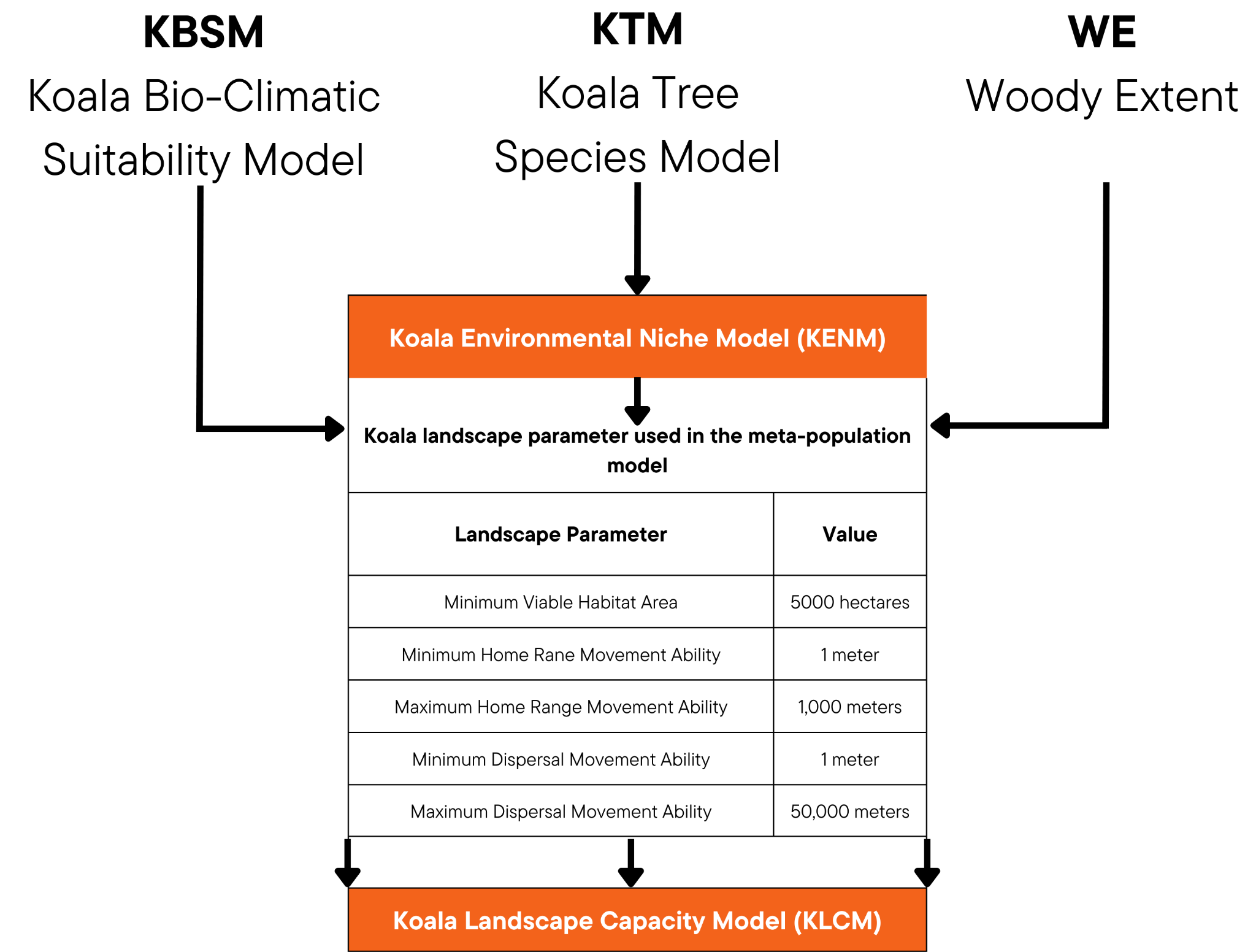

**Figure S2.1.** Methodological flow of Koala Landscape Capacity Model

REMP is a mechanistic model which considers the amount of habitat available to support populations and the habitat connectivity available to support dispersals to vacant habitats. Its output evaluates each location in a study in terms of capacity to support populations. This requires a modest set of species-specific landscape characteristics (or parameters, shown in the large box in Figure S2.1). The model was run over a seven epoch time-series (decadal between 2000 - 2070) for each combination of four NARCliM v.1 General Circulation Models (CSIRO-Mk3.0, ECHAM5, MIROC3.2, and CCMA 3.1) and three Regional Climate scenarios (which cover a limited area of the globe and are run at much finer spatial resolution compared to global model) (Evans et al. 2014). The KTM and KBSM models were linearly interpolated for each decade between the baseline (2000), 2030 and 2070 as part of the process of deriving a time-series of KENMs and ultimately KLCMs. For this analysis we used the baseline (2000) and 2070 (based on an averaged KENM across GCMs) time points to represent current and future conditions, respectively. The predictions are for each 90m by 90m grid cells across NSW.

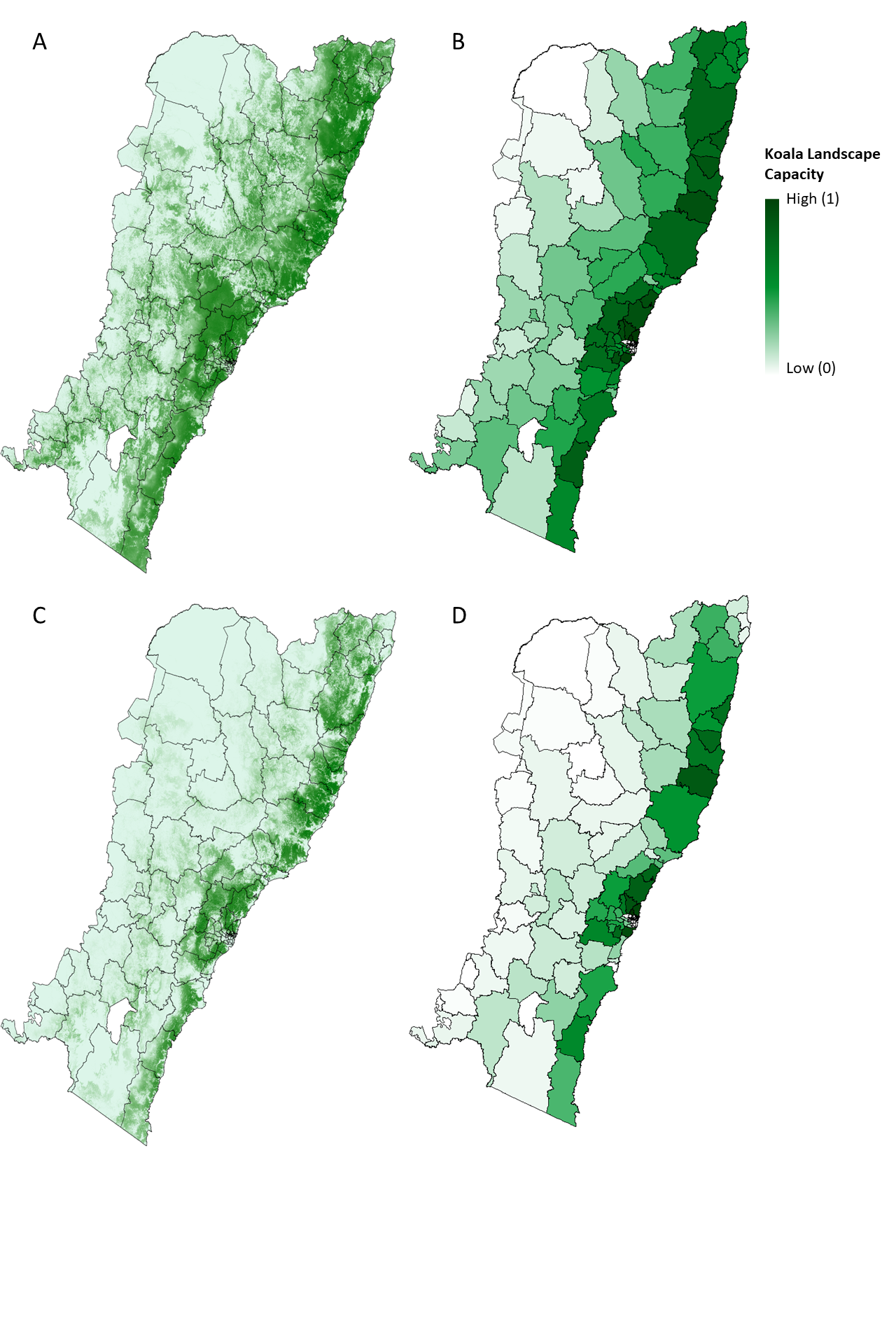

**Figure S2.2.** Koala landscape capacity under A) current climatic conditions, B) aggregated by mean to the LGA level, and C) future climatic conditions (2070), D) aggregated to by mean the LGA level

#### Supporting Material 3 - Landholder preference models

##### 3.1 Landholder survey

As part of a broader project on landholder engagement in private land conservation in the study area we developed and implemented an online survey instrument (on the Qualtrics platform) to ask landholders about various aspects of their conservation activities, their livelihoods as it relates to land-use, their willingness to participate in conservation covenants, and how impacted they are by climate change and extreme weather events. This survey complied with the requirements of the Human Research Ethics Committee from the University of Queensland (Human Ethics Approval #2019002363).

We implemented the survey across the study area using three different recruitment strategies. First, participants were recruited through a social research company (PureProfile) using screening criteria which screened out landholders who were not in New South Wales and did not own or manage land in a rural area greater than 2 hectares in size. These participants were financially compensated through PureProfile upon survey completion. Second, we recruited participants through a mail-out (containing a link to the online survey) to landholders addressed to a randomly stratified selection of 20,000 private properties greater than 2 hectares in size across the study area. Addresses were obtained from the information services company Experian and stratified according to postcode and Experian’s social segments (<http://www.experian.com.au/mosaic>) to ensure demographic representativeness. Mail-out participants could choose between two options to be compensated: either financially (a $10 Coles Myer voucher) or by requesting a $10 donation to a charity (Australian Foodbank, Bush Heritage Australia, or Australian Red Cross). Finally, we recruited participants by email through government organisational networks that administer programs using a snowball sampling technique (Johnson 2014).

Within the survey, landholders were asked to identify the financial payment they would be willing to accept to agree to a conservation covenant. We presented participants with a hypothetical conservation covenant program and asked participants to state how much money per hectare per year would be the minimum amount they would be willing to accept (willingness to accept) to join this program for both an in-perpetuity duration covenant and a 10 year duration covenant. Payment options ranged from, “I would pay”, $0, $25, $50, $100, $250, $500, $750, $1,000, $1,500, $2,000, $2,500, >$2,500 and “I would not participate”. We also asked landholders to state what proportion of their property they would be prepared to covenant for that payment. Property locations were obtained either from the respondents’ stated address or the property closest to the stated nearest street corner or based on the mail-out address.

Prior to analysis we screened the responses to ensure only completed surveys were included, that respondents were either the landholder or land manager, and removed responses that took less than 5 minutes to complete to ensure we included responses with a minimum quality level. For statistical analysis we only considered responses where we had location information for the property (in cases where only the nearest street corner was provided by the respondent, we used the nearest property to the street corner) and excluded properties that had intensive land-uses on them (specifically these were properties that had a secondary land use of “Manufacturing and Industrial” or “Services”, or a tertiary land use of “Urban Residential” based on NSW Government land-use mapping - <https://datasets.seed.nsw.gov.au/dataset/nsw-landuse-2017-v1p2-f0ed>). This resulted in 311 valid landholder responses (67 from PureProfile, 29 from the snowball sampling, and 215 from the mail-out survey).

##### 3.2 Statistical models

Based on the survey data we developed Bayesian statistical regression models of three landholder preference quantities: (1) the probability a landholder would be willing to consider participating in a conservation covenant (based the “I would not participate” response), (2) the minimum payment per hectare per year a landholder would be willing to accept in return for participating in a conservation covenant on their property (willingness to accept), and (3) the proportion of their property a landholder would be willing to covenant. We used statements about whether landholders would consider participating in a covenant or not as a proxy for their preference to put in an expression of interest during a tender process (SM 1.3). We built three different regression models for each quantity based on the following distributional assumptions: (1) landholder statements about whether they would consider a covenant or not had a Bernoulli distributed (i.e., logistic regression), (2) landholder statements about the payments they would be willing to accept to participate in a covenant had a normal distribution with appropriate censoring to account for the ranges defined in the survey question (i.e., linear regression), and (3) landholder statements about the proportion of their property they would covenant had a beta distribution (i.e., beta regression).

We developed a range of spatial explanatory variables that could be measured for all properties in the study area based on variables that were hypothesised to potentially be important for determining each of the three quantities modelled (see Table S3.2.1 for a list and justification for each variable). Depending on the resolution of data available, explanatory variables were characterised either at the property resolution or at the 2016 Australian Census Statistical Area 1 (SA1) spatial unit resolution (areas containing around 200-800 people [<https://www.abs.gov.au/statistics/standards/australian-statistical-geography-standard-asgs-edition-3/jul2021-jun2026/main-structure-and-greater-capital-city-statistical-areas/statistical-area-level-1#sa1-design-criteria>]). We also included variables related to the geographical context of each property, such as the distance to urban centres and the Koala Modelling Region (regions defined by the New South Wales Government for modelling koala populations) within which a property was located. Finally, we included SA1 as an intercept random-effect in the models to control for unexplained variation among SA1s and to account for ant spatial autocorrelation present among properties within the same SA1.

Although the primary aim of the model was prediction, rather than inference about the effect of each explanatory variable, we still aimed to partially control for collinearity among explanatory variables to ensure we got stable parameter estimates. Prior to model fitting we assessed the continuous explanatory variables measured at the property resolution for collinearity and where the correlation coefficient between two variables was > 0.6 we excluded one of the variables. This resulted in the exclusion of property size, distance to major urban centres, proportion of property native tree cover, ecological connectivity, and Terrain Ruggedness Index (Table 1). Principal components derived from the demographic variables measured at the SA1 resolution were all retained regardless of collinearity with the variables measured at the property resolution, so as to ensure we maintained predictors measured at the SA1 resolution. Similarly, all categorical explanatory variables (Koala Modelling Region, Mosaic Social Segment type, and Land-use) were retained despite evidence of collinearity because each variable was chosen to represent a specific distinct characteristic. Missing data in the explanatory variables were imputed using the R package “mice” with 10 replicate sets of imputed values generated (Azur et al. 2011; van Buuren & Groothuis-Oudshoorn 2011). Note that where the location information available was only either a road name or nearest street corner the property level covariates were set to missing data and imputed, while SA1 level covariates were retained for these properties. Separate regression models were developed for 10-year and in-perpetuity covenant survey responses although only the in-perpetuity models were used for this study.

Models were fit to the data via Markov Chain Monte Carlo (MCMC) with uninformative priors using JAGS (<https://mcmc-jags.sourceforge.io/>) and the R package “rjags”. We used three MCMC chains with overdispersed starting values, a burnin of 10,000 iterations and then retaining the next 10,000 iterations. Convergence was assessed using the Gelman-Rubin statistic (Gelman & Rubin 1992). To assess the importance of each explanatory variable in each model we assessed variable selection probabilities using the method described by Chen et al. (2016). This approach provides a way to assess the probability that a variable should be in the model and provide model-averaged coefficient estimates so that coefficients for explanatory variables with low inclusion probabilities are shrunk towards zero. Finally, we used the model-averaged coefficient estimates to make predictions of each landholder preference quantity for all properties > 2 ha in size with non-intensive land uses in the study area (Figure S3.2.1). All code for fitting the models and generating the predictions are available at: <https://github.com/koala-private-land/spatial-bid-model>.

**Table S3.2.1.** Explanatory variables included in the regression models

| Variable Type | Explanatory Variables | Data Source | Rationale and Notes |
| --- | --- | --- | --- |
| Demographics (Statistical Area 1 resolution) | Age, proportion Year 12 educated, proportion Bachelor Degree educated, proportion born in Australia, proportion English spoken at home, proportion parents’ born overseas, mean household size, proportion of four household composition categories, mean household income, mean mortgage repayments, proportion employed in agriculture (numeric). | Australian Bureau of Statistics 2016 Census (<https://www.abs.gov.au/websitedbs/censushome.nsf/home/2016>). | Demographics have been shown to be important predictors of landholder participation in private land conservation programs (Simmons et al. 2020). For example, older landholders might be more inclined to consider in-perpetuity covenants due to a longer-term perspective on property management and a desire to leave a lasting legacy for future generations. Similarly, landholders with higher education levels might be more aware of environmental and conservation concerns, making them more likely to consider such covenants that promote sustainable land use. Higher income landholders might be more willing to invest in conservation efforts, whereas lower income landholders might be more concerned about immediate financial needs and less likely to consider long-term commitments.  Principal components analysis was used to reduce dimensions across demographic variables and the first five principal components were used as explanatory variables. |
| Demographics (property resolution) | Mosaic Social Segment types (categorical). | Experian (<http://www.experian.com.au/mosaic>). | Rationale as above.  These are demographic categories developed by the market research company Experian. The following categories (with at least five responses in each category) were used: D13 (retired, traditional couples living in coastal and scenic areas, with average pensionable income levels, E16 (working in trades, middle-aged families owning acreages of land with large properties just outside the metro fringe), N48N49 - N48 (rural farmers and farm owners with below average income, living 10-40 km away from the nearest town) & N49 (very rural farmers and farm owners with below average income, living > 40 km from the nearest town), N50N51 - N50 (single farm workers in very small rural towns. with low income and low value properties) & N51 (low education, traditional, singles in far inland remote towns, with low income and low value properties), OTHER (all other types). |
| Economic (property resolution) | Unimproved land value (numeric, $/ha). | NSW Land Value Information 2016 (<https://data.nsw.gov.au/data/dataset/http-www-valuergeneral-nsw-gov-au-land-value-summaries-lv-php>). | Land value might influence people's preferences for private land conservation programs. For example, landholders with lower unimproved land values might be more sensitive to the opportunity costs of participating in conservation programs. If the land has a lower potential for development or income generation, these landholders might be more likely to consider conservation to maximise the land's value.  Log(unimproved land value) was used as the explanatory variable. |
|  | Dominant (most common) land-use (categorical). | NSW Landuse 2017 v1.2 (<https://datasets.seed.nsw.gov.au/dataset/nsw-landuse-2017-v1p2-f0ed>). | In areas where the dominant land use is heavily developed, people might be more attuned to the potential conflicts between development and conservation. This awareness could lead to increased support for initiatives that protect natural areas. They may also recognise the economic importance of conservation to sustain their livelihoods.  The following categories (five most common responses with at least five responses in each category) were used: 1.3.0 Other minimal use, 2.1.0 Grazing native vegetation, 3.2.0 Grazing modified pastures, 5.4.0 Residential and farm infrastructure, 3.3.0 cropping, OTHER. Codes refer to the Australian Land Use and Management (ALUM) Classification codes. |
|  | Property size (numeric, ha) | NSW Land Parcel and Property Theme (<https://portal.spatial.nsw.gov.au/portal/home/item.html?id=01de8834e88a45a1a673b120aa00c82e>). | Larger properties might have more potential for diverse use. Larger properties often have more space to accommodate diverse land uses. Owners might feel more comfortable dedicating a portion of their land to conservation without significantly affecting other potential uses.  Log(property size) was used as the explanatory variable. |
|  | Dominant (most common) soil capability value (numeric). | Land and Soil Capability Mapping for NSW (<https://datasets.seed.nsw.gov.au/dataset/land-and-soil-capability-mapping-for-nsw4bc12>). | Landowners with soils more suitable for agricultural production might be more inclined to use their land for farming. Participating in a conservation program might be seen as a way to balance conservation goals with agricultural productivity.  Values used are based on an eight class system with values ranging between 1 and 8 which represent a decreasing capability of the land to sustain land-use. Class 1 represents land capable of sustaining most land-uses including those that have a high impact on the soil (e.g., regular cultivation), whilst class 8 represents land that can only sustain very low impact land-uses (e.g., nature conservation). This index is treated as numeric in the regression models. |
| Geographic Context | Distance to Major Urban Centres (numeric, km) | Australian Statistical Geography Standard Section of State Volume 4 (<https://www.abs.gov.au/ausstats/abs@.nsf/Lookup/by%20Subject/1270.0.55.004~July%202016~Main%20Features~Design%20of%20SOS%20and%20SOSR~12>). | Landowners closer to urban centres might prioritise commercial activities due to easier access to markets, making them more hesitant to allocate land to conservation if they perceive it as limiting economic opportunities.  Log(distance to major urban centres + 1) was used as the explanatory variable. |
|  | Distance to Other Urban Centres (numeric, km) | Australian Statistical Geography Standard Section of State Volume 4 (<https://www.abs.gov.au/ausstats/abs@.nsf/Lookup/by%20Subject/1270.0.55.004~July%202016~Main%20Features~Design%20of%20SOS%20and%20SOSR~12>). | Same as previous.  Log(distance to other urban centres + 1) was used as the explanatory variable. |
|  | Koala Modelling Region (categorical). | NSW Department of Planning and Environment (<https://datasets.seed.nsw.gov.au/dataset/koala-modelling-regions>). | Differences in how koalas and other biodiversity are managed across different regions might influence landholders values for conservation. |
| Ecological (property resolution) | Proportion of property native tree cover (numeric). | NSW Native Vegetation Extent 5m Raster v1.2 (<https://datasets.seed.nsw.gov.au/dataset/nsw-native-vegetation-extent-5m-raster-v1-0>). | Landowners with high native tree cover might be more motivated to participate in conservation programs due to the potential for supporting diverse ecosystems and wildlife habitats. |
|  | Proportion of property native grass cover (numeric). | NSW Native Vegetation Extent 5m Raster v1.2 (<https://datasets.seed.nsw.gov.au/dataset/nsw-native-vegetation-extent-5m-raster-v1-0>). | Same as previous. |
|  | Ecological condition (numeric). | Ecological condition of terrestrial habitat (<https://datasets.seed.nsw.gov.au/dataset/ecological-condition-of-terrestrial-habitat>). | Same as previous. |
|  | Ecological connectivity (numeric). | Ecological connectivity of terrestrial habitat (<https://datasets.seed.nsw.gov.au/dataset/ecological-connectivity-of-terrestrial-habitat>). | Same as previous. |
| Biophysical (property resolution) | Elevation (numeric, m) | Geosciences Australia 1 Second Digital Elevation Model, DEM-S (<https://ecat.ga.gov.au/geonetwork/srv/eng/catalog.search#/metadata/72759>). | Higher elevations often host unique ecosystems and species adapted to cooler temperatures. Landowners might be motivated to participate in conservation programs to protect these specialised habitats.  Log(elevation) was used as the explanatory variable. |
|  | Slope (numeric, degrees) | Geosciences Australia 1 Second Digital Elevation Model, DEM-S (<https://ecat.ga.gov.au/geonetwork/srv/eng/catalog.search#/metadata/72759>). | Properties with easy access are more convenient to manage and monitor. Landowners might be more willing to participate in conservation programs if there is less slope and they can easily oversee and maintain the required practices.  Slope was calculated using ArcGIS Pro Version 3.0.2. |
|  | Terrain Ruggedness Index (numeric) | Geosciences Australia 1 Second Digital Elevation Model, DEM-S (<https://ecat.ga.gov.au/geonetwork/srv/eng/catalog.search#/metadata/72759>). | Areas with low ruggedness might be more suitable for agriculture due to ease of cultivation and access. Landowners might be less inclined to participate if conservation limits agricultural potential.  Terrain Ruggedness Index is an index of the difference in elevation between adjacent raster cells. Terrain Ruggedness Index was calculated using ArcGIS Pro Version 3.0.2. |

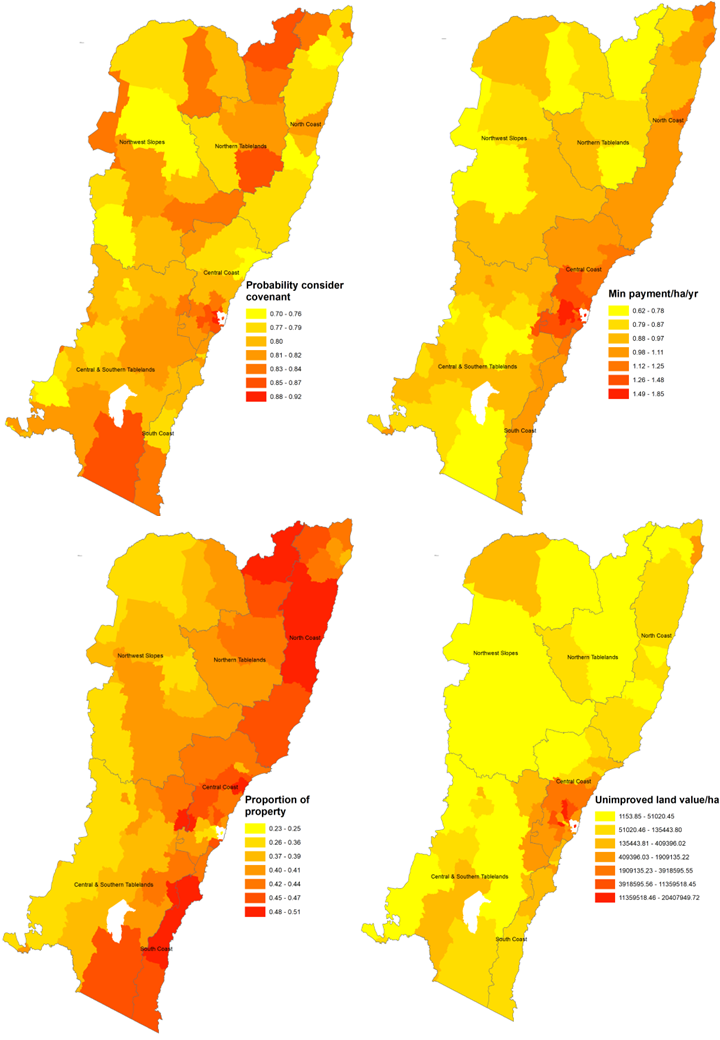

**Figure S3.2.1.** Mean spatial predictions for private properties containing koala habitat for each LGA for: (A) the probability a landholder would consider an in-perpetuity covenant, (B) the stated financial payment required to adopt an in-perpetuity covenant, and (C) the stated proportion of their property a landholder would consider applying an in-perpetuity covenant to. (D) shows the mean unimproved land value per ha $AUD (New South Wales Government 2017). Data has been aggregated by mean to the LGA level.

#### Supporting Material 4 - Cost efficiency

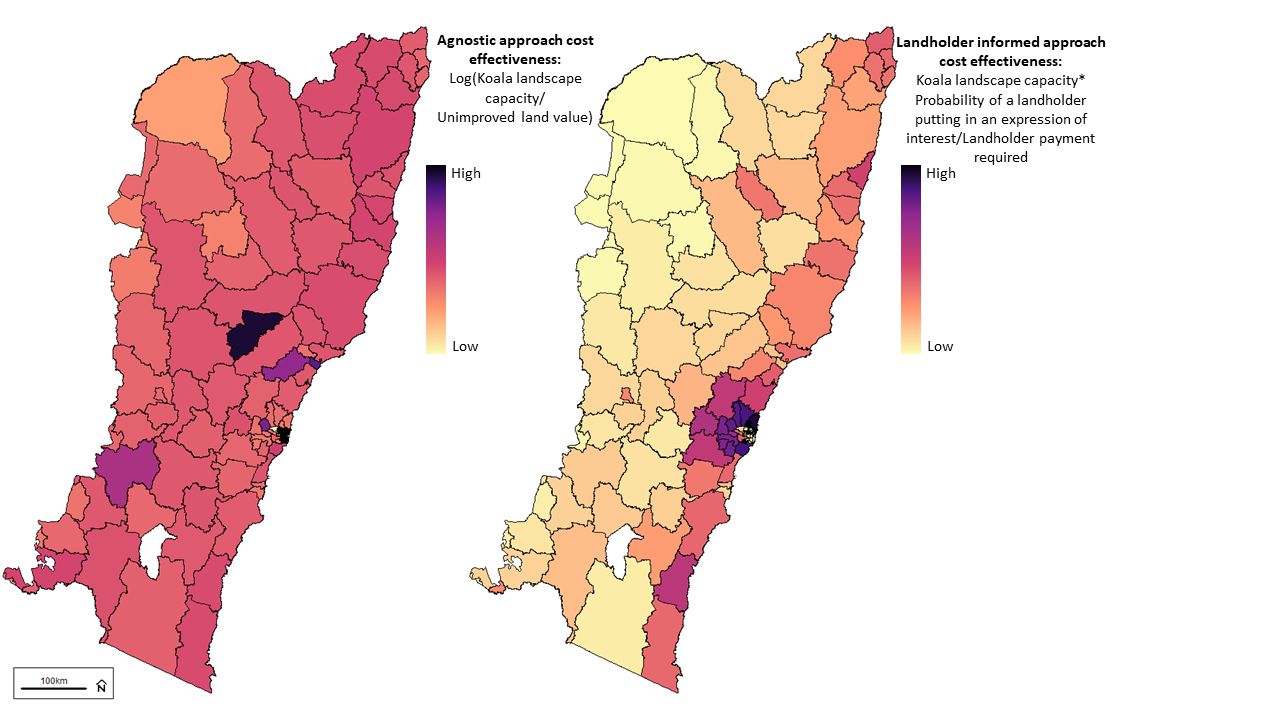

**Figure S4.1** - Supporting figures showing the cost efficiency of investing in properties, the mean values aggregated at the LGA scale, for the agnostic approach (left) and the landholder informed approach (right)

#### Supporting Material 5 - Values and uncertainty for all approaches and scenarios

| Approach | Budget  (million $AUD) | Cost  (million $AUD) | Conservation  benefit | Summed probability  of a bid | SD | SE |
| --- | --- | --- | --- | --- | --- | --- |
| Agnostic | 20.3 | 19.307 | 4.071 | 30.507 | 0.197 | 0.014 |
| Agnostic | 40.6 | 39.994 | 4.129 | 35.562 | 0.146 | 0.01 |
| Agnostic | 60.9 | 60.333 | 4.369 | 39.267 | 0.127 | 0.009 |
| Agnostic | 81.2 | 80.847 | 7.696 | 68.329 | 0.13 | 0.011 |
| Agnostic | 101.5 | 101.36 | 9.953 | 87.69 | 0.135 | 0.011 |
| Agnostic | 121.8 | 121.699 | 12.868 | 97.978 | 0.178 | 0.014 |
| Agnostic | 142.1 | 141.689 | 12.547 | 102.651 | 0.17 | 0.013 |
| Agnostic | 162.4 | 162.028 | 14.364 | 108.966 | 0.171 | 0.013 |
| Agnostic | 182.7 | 182.192 | 14.853 | 117.947 | 0.165 | 0.013 |
| Agnostic | 203 | 202.531 | 15.211 | 120.85 | 0.16 | 0.012 |
| Landholder informed | 20.3 | 20.084 | 25.097 | 79.772 | 0.295 | 0.021 |
| Landholder informed | 40.6 | 40.598 | 32.806 | 106.141 | 0.275 | 0.019 |
| Landholder informed | 60.9 | 60.333 | 39.237 | 129.729 | 0.283 | 0.021 |
| Landholder informed | 81.2 | 80.498 | 43.752 | 150.309 | 0.299 | 0.021 |
| Landholder informed | 101.5 | 101.44 | 43.538 | 177.382 | 0.334 | 0.024 |
| Landholder informed | 121.8 | 121.779 | 46.467 | 197.342 | 0.281 | 0.02 |
| Landholder informed | 142.1 | 141.943 | 48.554 | 212.256 | 0.285 | 0.02 |
| Landholder informed | 162.4 | 162.282 | 50.842 | 232.444 | 0.266 | 0.019 |
| Landholder informed | 182.7 | 182.621 | 52.345 | 254.522 | 0.25 | 0.018 |
| Landholder informed | 203 | 202.96 | 54.593 | 280.918 | 0.248 | 0.018 |
| Climate change informed | 20.3 | 20.084 | 3.67 | 19.146 | 0.231 | 0.016 |
| Climate change informed | 40.6 | 39.82 | 3.978 | 24.18 | 0.162 | 0.012 |
| Climate change informed | 60.9 | 60.333 | 7.107 | 50.932 | 0.14 | 0.01 |
| Climate change informed | 81.2 | 80.847 | 7.696 | 55.63 | 0.114 | 0.008 |
| Climate change informed | 101.5 | 101.185 | 9.388 | 64.399 | 0.141 | 0.01 |
| Climate change informed | 121.8 | 121.35 | 9.375 | 74.805 | 0.106 | 0.008 |
| Climate change informed | 142.1 | 141.689 | 13.776 | 88.929 | 0.192 | 0.014 |
| Climate change informed | 162.4 | 161.679 | 13.712 | 110.25 | 0.202 | 0.014 |
| Climate change informed | 182.7 | 181.844 | 13.926 | 113.354 | 0.182 | 0.013 |
| Climate change informed | 203 | 202.96 | 12.958 | 110.223 | 0.145 | 0.01 |
| Integrated | 20.3 | 19.91 | 24.123 | 78.654 | 0.234 | 0.017 |
| Integrated | 40.6 | 40.598 | 33.367 | 104.846 | 0.258 | 0.018 |
| Integrated | 60.9 | 60.159 | 39.165 | 125.315 | 0.229 | 0.016 |
| Integrated | 81.2 | 80.982 | 43.324 | 139.327 | 0.203 | 0.015 |
| Integrated | 101.5 | 101.32 | 47.088 | 161.795 | 0.201 | 0.014 |
| Integrated | 121.8 | 121.35 | 50.74 | 172.294 | 0.205 | 0.015 |
| Integrated | 142.1 | 141.943 | 54.213 | 181.81 | 0.247 | 0.017 |
| Integrated | 162.4 | 162.282 | 57.567 | 193.113 | 0.243 | 0.017 |
| Integrated | 182.7 | 182.621 | 60.648 | 206.733 | 0.224 | 0.016 |
| Integrated | 203 | 202.921 | 63.779 | 212.259 | 0.269 | 0.019 |

#### Supporting Material 6 - The Biodiversity Conservation Trust’s Biodiversity Valuation Score and the data that goes into it.

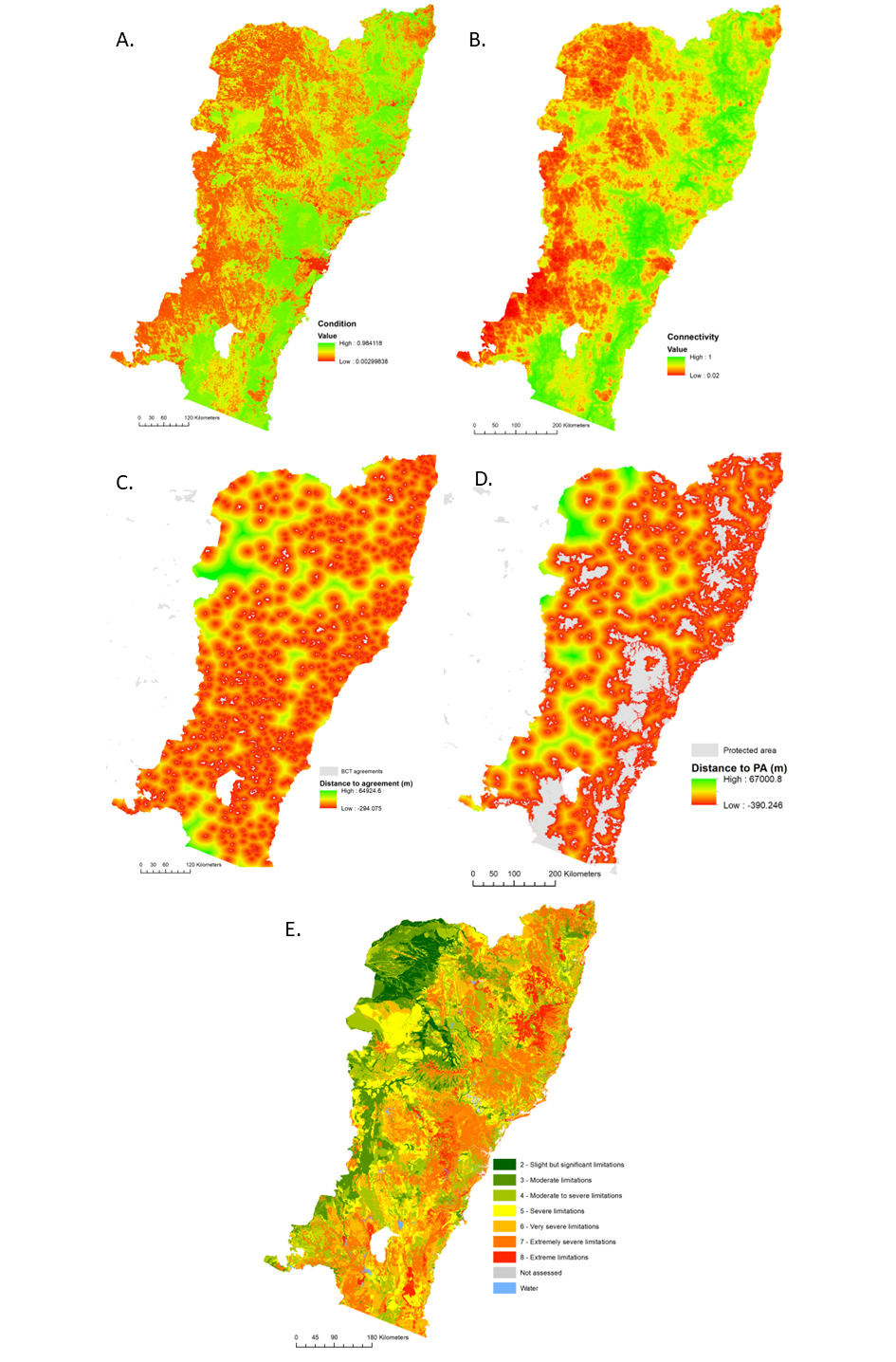

**Figure S6.1** - Input data used to calculate the Biodiversity Conservation Trust’s Assessment Metric. Datasets were A) Condition (Love et al. 2018), B) Connectivity (Love et al. 2018), C) Distance to other conservation agreements, D) Distance to protected areas, and E) Risk of habitat conversion derived from land and soil capability as a proxy (DPE 2021).

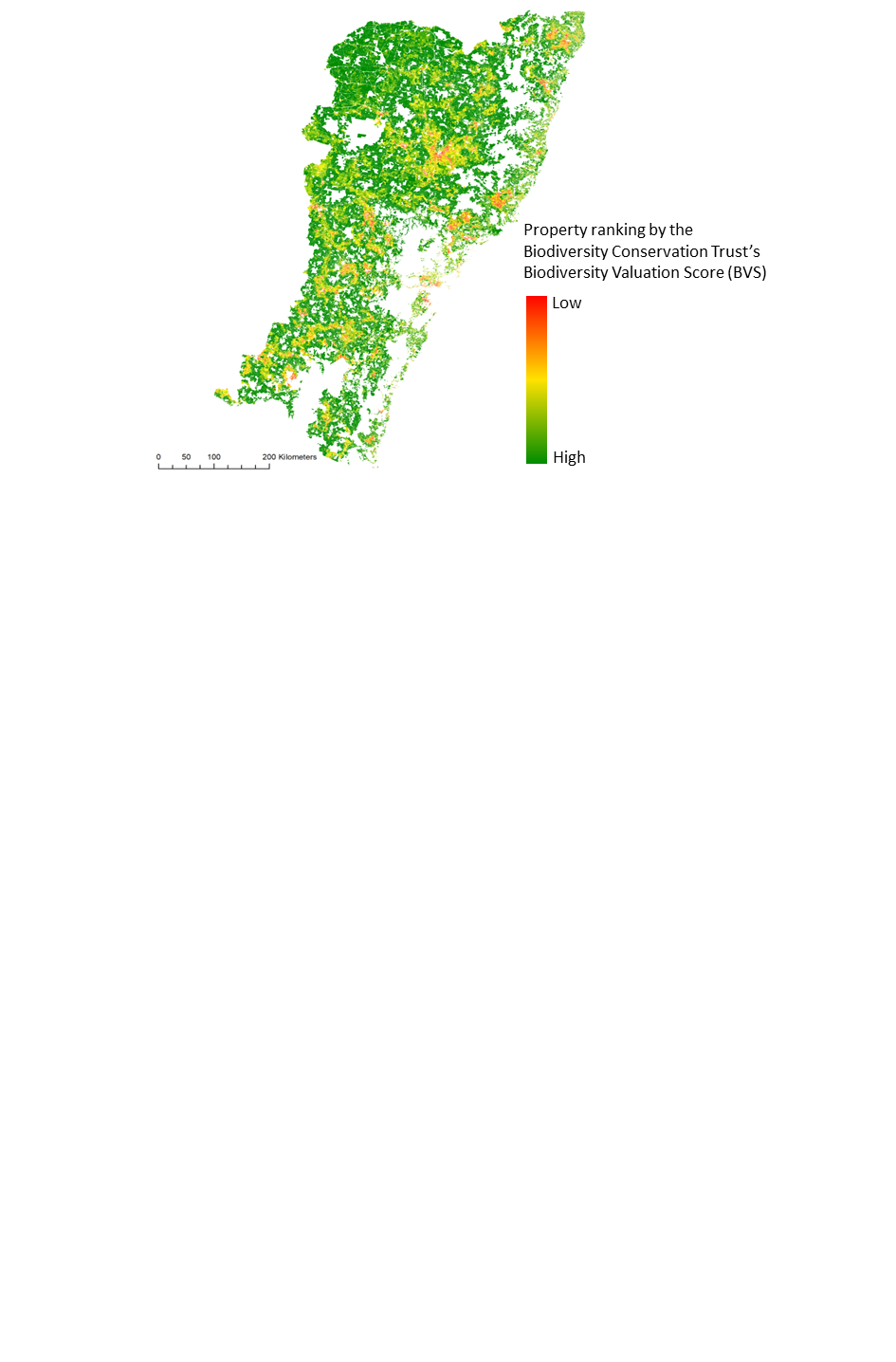

**Figure S6.2** - Properties ranked by the Biodiversity Conservation Trust’s Biodiversity Valuation Score (BVS)

#### Supporting Material 7 - Informing on NSW Koala Strategy Objectives

##### 7.1 Scenarios to inform on NSW Koala Strategy Objectives

In addition to exploring a suite of budget scenarios, we explored a further scenario to inform on the NSW Koala Strategy’s objectives. We used the 19 priority populations for immediate investment identified in the NSW Koala Strategy as koala populations for immediate investments (NSW Government 2022c) as planning units and identified optimal investments under a $20.3 million overall budget using integrated planning approach.

##### 7.2 Findings to inform on NSW Koala Strategy Objectives

###### 7.2.1 Budget comparison

We find a budget of $20.3 M, the budget announced in the 2022 koala strategy to protect koala habitat (over the next 5 years) on private land, could result in the protection through tenders of 4.49 km^2^ of koala habitat, $60.9 million could result in 16.4 km^2^, $101.5 M in 28.7 km^2^, $142.1 M in 46.3 km^2^, $162.4 M in 48.8 km^2^, and $203 M in 62.8 km^2^ using the integrated approach across the entire study region (SM5).

###### 7.2.2 NSW Koala Strategy Priority Populations

Of the 19 priority areas earmarked for priority investment Armidale was allocated $10.8M, Whiporie - Rappville $5.9M, and the Blue Mountains $3.03M of the $20.3 M budget using as the overall budget in integrated approach. This resulted in 6.20 km^2^ of koala habitat with high koala landscape capacity conserved (Figure S7.2.2.1).

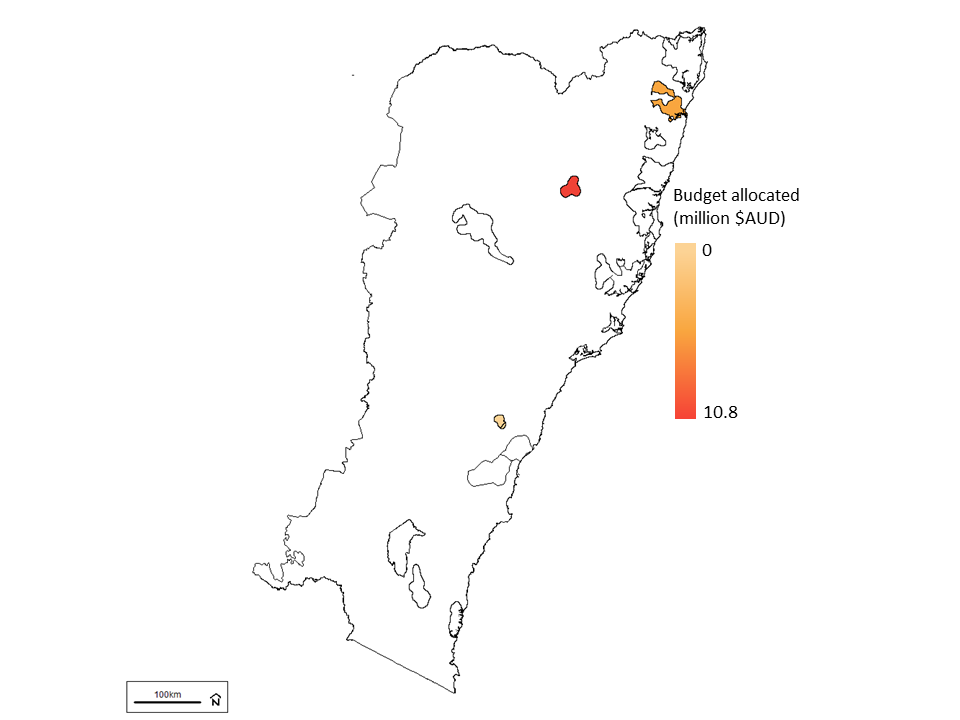

**Figure S7.2.2.1** - Budget allocations under the integrated approach (#4 - which considers landholder preferences and future climatic conditions) with an overall budget of $20.3 M AUD (the amount of the 2022 NSW Koala Strategy's budget allocated to protecting koala habitat on private land through Biodiversity Conservation Trust’s Conservation Partners Program (not considered in this paper) and in-perpetuity conservation agreements with annual payments to private landholders) within the 19 priority areas earmarked for priority investment. This resulted in 6.20 km^2^ of conserved areas with high koala landscape capacity.

### References

Azur, M.J., Stuart, E.A., Frangakis, C. & Leaf, P.J. (2011). Multiple imputation by chained equations: what is it and how does it work? *Int. J. Methods Psychiatr. Res.*, 20, 40–49.

Biodiversity Conservation Trust. (2020). Biodiversity Conservation Trust Assessment Metric.

van Buuren, S. & Groothuis-Oudshoorn, K. (2011). mice: Multivariate imputation by chained equations in R. *J. Stat. Softw.*, 45, 1–67.

Chen, R.-B., Chu, C.-H., Yuan, S. & Wu, Y.N. (2016). Bayesian sparse group selection. *J. Comput. Graph. Stat.*, 25, 665–683.

DPE. (2021). Land and Soil Capability Mapping for NSW, Version 4.5. *NSW Department of Planning, Industry and Environment, Parramatta*. <https://datasets.seed.nsw.gov.au/dataset/land-and-soil-capability-mapping-for-nsw4bc12>

Drielsma, M., Love, J., Thapa, R., Thonell, J. & Beaumont, L. (In Prep). Koalas in the landscape (KITL) - Landscape capacity to support Koala populations through climate change. *Department of Planning, Industry, and Environment, NSW Government*.

Evans, J.P., Argüeso, D. & Di Luca, A. (2014). Design of a Regional Climate Model Ensemble That Incorporates Model Performance and Independence. *AGU Fall Meeting*. <https://ui.adsabs.harvard.edu/abs/2014AGUFM.A53Q..01E/abstract>

Gelman, A. & Rubin, D.B. (1992). Inference from iterative simulation using multiple sequences. *Stat. Sci.*, 7, 457–472.

Hajkowicz, S., Higgins, A., Miller, C. & Marinoni, O. (2008). Targeting conservation payments to achieve multiple outcomes. *Biol. Conserv.*, 141, 2368–2375.

Hajkowicz, S., Higgins, A., Williams, K., Faith, D.P. & Burton, M. (2007). Optimisation and the selection of conservation contracts. *Aust. J. Agric. Resour. Econ.*, 51, 39–56.

Johnson, T.P. (2014). Snowball Sampling: Introduction. *Wiley StatsRef: Statistics Reference Online*.

Love, J., Drielsma, M.J., Williams, K. & Thapa, R. (2018). Data package for habitat condition indicators; 3.1a ecological condition, 3.1b ecological connectivity and 3.1c ecological carrying capacity. *NSW Office of Environment and Heritage*. <https://datasets.seed.nsw.gov.au/dataset/2cf9b633-1b4e-43a0-a363-477c5bc08988>

Marinoni, O., Higgins, A. & Hajkowicz, S. (2008). Development of a natural resource management investment decision support system. *Commonwealth Scientific and Industrial Research Organisation (CSIRO) Sustainable Ecosystem*. <https://www.researchgate.net/profile/Oswald-Marinoni/publication/228494558_Development_of_a_Natural_Resource_Management_Investment_Decision_Support_System/links/0c9605255f5061221a000000/Development-of-a-Natural-Resource-Management-Investment-Decision-Support-System.pdf>

Marinoni, O., Higgins, A., Hajkowicz, S. & Collins, K. (2009). The multiple criteria analysis tool (MCAT): A new software tool to support environmental investment decision making. *Environmental Modelling & Software*, 24, 153–164.

New South Wales Government. (2017). Bulk land value information available on the Valuation Services Portal. <https://data.nsw.gov.au/data/dataset/327f9982-e610-4ffe-a51c-4047e97c7e7d>

NSW Government. (2022a). NSW Land Parcel Property Theme. <https://portal.spatial.nsw.gov.au/server/rest/services/NSW_Land_Parcel_Property_Theme/FeatureServer>

NSW Government. (2022b). NSW National Parks and Wildlife Service (NPWS) Estate. <https://datasets.seed.nsw.gov.au/dataset/nsw-national-parks-and-wildlife-service-npws-estate3f9e7>

NSW Government. (2022c). NSW Koala Strategy - Towards doubling the number of koalas in New South Wales by 2050. <https://www.environment.nsw.gov.au/-/media/OEH/Corporate-Site/Documents/Animals-and-plants/Threatened-species/koala-strategy-2022-220075.pdf>

Simmons, B.A., Archibald, C.L., Wilson, K.A. & Dean, A.J. (2020). Program Awareness, Social Capital, and Perceptions of Trees Influence Participation in Private Land Conservation Programs in Queensland, Australia. *Environ. Manage.*, 66, 289–304.
